# Systematic Meta-Analysis of Published Transcriptomic Prognostic Signatures and Development of a Robust Multi-Cohort Prognostic Classifier for Triple-Negative Breast Cancer

**DOI:** 10.64898/2026.07.09.737472

**Authors:** Leepakshi Dhingra, Manulaya Singh, Simran Jit, Deepika Yadav, Sherry Bhalla

## Abstract

Triple-negative breast cancer (TNBC) exhibits pronounced molecular heterogeneity, yet the majority of published transcriptomic prognostic signatures suffer from limited reproducibility and have not achieved clinical translation. We systematically benchmarked 62 published TNBC prognostic signatures across 6 independent cohorts (n=1,357) using a unified analytical framework spanning multiple scoring algorithms, survival endpoints, and threshold strategies. While 17 signatures demonstrated consistent univariate prognostic associations, only 4 remained independently prognostic after adjustment for clinicopathological variables, and none achieved robustness across all analytical conditions-underscoring the fragility of existing classifiers.

Leveraging genes with concordant survival associations across all 6 discovery cohorts, we identified a reproducible 15-gene directionally concordant gene set (DCGS) signature and distilled it into MetaSig-EFS, a 13-gene prognostic model optimized using a cohort-aware DeepSurv framework. MetaSig-EFS demonstrated robust cross-cohort generalizability, achieving validation concordance indices of 0.89 in GSE19615 and 0.69 in JBordet, with corresponding 3- and 5-year time-dependent AUROCs of 0.89 and 0.96 in GSE19615 and 0.80 and 0.71 in JBordet, respectively, while retaining independent prognostic value across established TNBC molecular sub-typing systems. Leveraging the TAHOE-100M transcriptomic perturbation atlas, we systematically prioritized candidate therapeutics through large-scale drug repurposing, identifying Paclitaxel as the top-ranked compound, followed by Venetoclax and Tucatinib. Together, these findings provide biologically informed and clinically actionable therapeutic hypotheses that extend the translational utility of our validated prognostic framework for TNBC risk stratification.

## Introduction

Triple-negative breast cancer (TNBC) presents a fundamental clinical challenge: the same diagnosis encompasses patients whose outcomes diverge dramatically, from long-term remission following chemotherapy to rapid metastatic progression and early death, yet the tools routinely available to distinguish these trajectories remain inadequate. Defined by the absence of estrogen receptor, progesterone receptor, and HER2 expression, TNBC historically restricted systemic treatment to cytotoxic chemotherapy^1^. Although immune checkpoint inhibitors and PARP inhibitors have improved outcomes in biomarker-selected patient subgroups, overall event rates remain substantially higher than in other breast cancer subtypes, and the majority of patients who relapse do so within the first three years^2,3^. This clinical unpredictability, compounded by the absence of targetable oncogenic drivers in most cases, makes reliable prognostic stratification beyond standard clinicopathological parameters - lymph node status, tumor size, and grade - an urgent and unmet need.

Large-scale molecular profiling has established that TNBC is not a single disease but a collection of biologically distinct states. These range from highly proliferative basal-like tumors enriched for genomic instability and cell cycle dysregulation, to immunologically active subtypes with dense T-cell infiltration and interferon signaling activity, to mesenchymal subtypes characterized by stromal remodeling and epithelial-mesenchymal transition^4^. Each of these states carries distinct prognostic implications: immune-active tumors tend to respond better to immunotherapy and fare better under chemotherapy, while mesenchymal and androgen receptor-expressing tumors show lower chemo-sensitivity and worse long-term outcomes. Capitalizing on this heterogeneity, numerous gene expression-based prognostic signatures have been developed over the past decade, designed to capture the dominant biological programs - proliferative activity, immune infiltration, stromal remodeling, and lineage-specific transcriptional states - and translate them into individual patient risk estimates.

Despite their biological promise, the clinical utility of these signatures has been constrained by several persistent and interrelated limitations. First, most signatures were derived from overlapping discovery cohorts, most notably TCGA-BRCA, which contains a limited number of TNBC samples and over represents patients who received adjuvant rather than neoadjuvant treatment; consequently, reported prognostic associations may reflect cohort-specific characteristics rather than generalizable biological signal^5^. Second, substantial heterogeneity in gene selection criteria, risk scoring frameworks, and survival endpoints makes direct comparison between models methodologically unreliable. Third, rigorous multi-cohort validation against established clinical variables - essential for demonstrating independent prognostic value beyond what pathologists already know from lymph node status and tumor size alone - has rarely been performed in a standardized and reproducible manner^4^. A particularly informative observation is that while published TNBC signatures rarely share individual genes, they tend to converge on a smaller set of recurrent biological processes including immune signaling, tumor proliferation, and extracellular matrix remodeling. This disconnect between gene-level divergence and pathway-level convergence suggests that many signatures capture related transcriptional programs through different molecular proxies, and that the field has been generating variations on a limited number of underlying biological themes - without the cross-cohort infrastructure needed to determine which variations are robust.

To address these gaps systematically, we evaluated 62 published TNBC prognostic gene signatures across six independent transcriptomic cohorts comprising 1,357 primary tumors profiled by microarray or RNA sequencing. Using standardized scoring methods, rigorous quality control, and multivariable survival analyses, we assessed the reproducibility and independent prognostic value of each signature. Leveraging a multi-cohort gene discovery framework requiring consistent survival associations across all six cohorts, we derived MetaSig - a novel prognostic classifier trained using cohort-weighted penalized Cox regression and a DeepSurv deep learning model, and externally validated in two independent cohorts. Risk stratification by MetaSig was assessed within clinically and molecularly defined subgroups, its biological underpinnings characterized through single-cell transcriptomics and master regulator inference, and its therapeutic implications explored through drug repurposing against the Tahoe-100M perturbation atlas. Together, this study provides a systematic and reproducible framework for evaluating TNBC transcriptomic signatures, and translates the resulting biological insights into a validated risk stratification tool with direct therapeutic relevance.

## Results

### Widespread Cohort Dependence and Biological Convergence Across Published TNBC Prognostic Signatures

Systematic literature curation identified 62 published prognostic gene signatures developed for TNBC (Supplementary Table S1A). We subsequently examined the patient cohorts, survival endpoints, gene composition, and pathway-level convergence underlying these models to define the biological processes represented across signatures. (Supplementary Table S1A-B). Signature size varied substantially ranging from 3 to 45 genes, with ∼58% of signatures containing 5-8 genes (Figure 1A). To characterize the data resources supporting transcriptomics-based TNBC prognostic models, we analyzed the cohorts employed for signature derivation and independent validation. TCGA-BRCA^6^ served as the predominant discovery dataset, being used for signature development in 42 of the 62 published models (67.7%) (Figure 1A). METABRIC^7^ was the second most commonly used cohort (13/62 signatures) for development. In contrast, cohorts such as GSE58812^8^, GSE21653^9^, SCAN-B^10^, and others were used more frequently for validation than discovery (Figure 1A). We next examined the survival endpoints used across the 62 signatures. Overall survival (OS) was the most commonly used endpoint, while a smaller number of studies used event-based outcomes, including recurrence-free, metastasis-free, disease-free, or event-free survival (Figure 1A). Only a few studies evaluated both endpoint types. Together, these findings reveal considerable variation in endpoint selection and a strong dependence of TNBC prognostic research on a limited set of publicly available cohorts.

**Figure 1:**
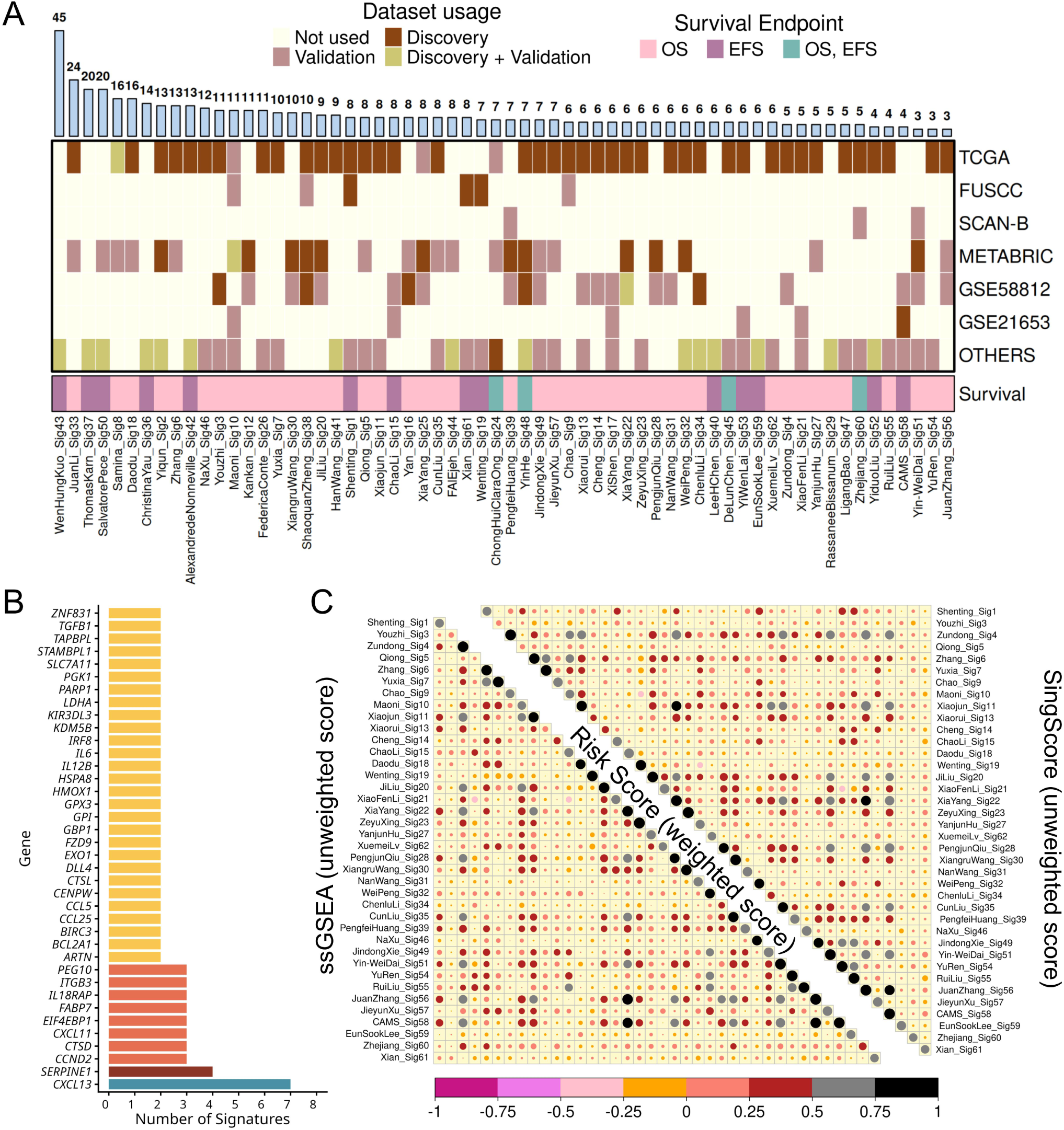
Characteristics of 62 published TNBC prognostic gene signatures. A) Distribution of signature length, discovery and validation datasets, and survival endpoints across 62 published signatures. B) Frequency of recurrent genes across signatures. C) Correlation of ssGSEA- and SingScore-derived scores with published coefficient-based RiskScores for 38 signatures with available formulas.

Despite the widespread reuse of two public cohorts, overlap at the individual gene level was limited. *CXCL13* was the most frequently represented gene, appearing in seven of the 62 signatures, followed by *SERPINE1*, which was present in four signatures. Eight additional genes - *CTSD, CCND2, CXCL11, EIF4EBP1, ITGB3, FABP7, PEG10*, and *IL18RAP* - were shared across three signatures each. A further 28 genes, including *BCL2A1, BIRC3, CCL25, CCL5, CTSL, DLL4, ARTN, CENPW, EXO1*, and *FZD9*, appeared in two signatures each (Figure 1B). Overall, only a small fraction of genes recurred across multiple signatures, with the vast majority appearing in a single model. Despite limited overlap at the gene level, Hallmark and C2^11^ pathway enrichment analyses revealed marked convergence on common biological programs, indicating that distinct TNBC prognostic signatures often capture similar underlying biology (Supplementary Figure S2A). Recurrently enriched pathway categories included immune regulation, stemness and developmental processes, oncogenic signaling, transcription factor networks, cell cycle and DNA repair, epigenetic and metabolic programs, stromal remodeling, and cellular stress responses (Supplementary Figure S2B; Supplementary Table S1C)

Among the 62 published TNBC prognostic signatures evaluated, only 38 provided explicit coefficient-based formulas for calculating patient-specific risk scores. We therefore evaluated whether coefficient-independent (unweighted) scoring approaches could provide a standardized framework for cross-signature comparison. To enable cross-cohort comparisons, signature scores were calculated using expression matrices that had been normalized and processed independently within each of the six discovery cohorts as described in the Methods section “Patient Cohorts and Data Processing” (Supplementary Figure S3A-H). The clinical and pathological characteristics of the 1,357 TNBC patients included across the six cohorts are summarized in Table 1. Each signature was scored using unweighted single sample scoring methods: scored using ssGSEA^12^ and SingScore^13^ which were Z-score normalized within each cohort (Supplementary Figure S4A-D). For the 38 signatures with available published formulas, pairwise correlation analysis showed strong concordance between the original weighted risk scores and the corresponding unweighted ssGSEA- and SingScore-derived scores (Figure 1C). The majority of signatures showed high positive correlations with the published weighted Risk Scores, with 30/38 (78.94%) of ssGSEA-derived scores achieving r > 0.7. SingScore showed a broadly comparable pattern of concordance with the original weighted models. These results indicate that patient ranking was largely preserved even when signature-specific coefficients were omitted, supporting the use of unweighted scoring methods for the systematic evaluation of all 62 TNBC prognostic signatures across independent cohorts

**Table 1.** Patient Characteristics in Discovery Cohorts.

| **Characteristics** | **GSE58812** | **GSE21653** | **METABRIC** | **SCAN-B** | **FUSCC** | **TCGA** | **TOTAL** |
| --- | --- | --- | --- | --- | --- | --- | --- |
| **TNBCs** | 107 | 87 | 320 | 326 | 360 | 157 | 1357 |
| **EFS STATUS**<br>**(Total)** | **107** | **85** | **320** | **320** | **360** | **141** | **1333** |
| 0 (no event) | 76 | 58 | 189 | 259 | 311 | 118 | 1011 |
| 1 (event) | 31 | 27 | 131 | 61 | 49 | 23 | 322 |
| NA | 0 | 2 | 0 | 6 | 0 | 16 | 18 |
| **OS STATUS**<br>**(Total)** | **107** | **-** | **320** | **326** | **258** | **152** | **1163** |
| 0 (alive) | 78 | - | 152 | 223 | 228 | 127 | 808 |
| 1 (dead) | 29 | - | 168 | 103 | 30 | 25 | 355 |
| NA | 0 | 87 | 0 | 0 | 102 | 5 | 194 |
| **LN STATUS**<br>**(Total)** | **-** | **85** | **299** | **321** | **358** | **144** | **1122** |
| 0 (negative) | - | 51 | 144 | 217 | 215 | 99 | 726 |
| 1 (positive) | - | 34 | 155 | 104 | 143 | 45 | 481 |
| NA | 107 | 2 | 21 | 5 | 2 | 13 | 150 |
| **SIZE**<br>**(Total)** | **-** | **85** | **312** | **316** | **359** | **157** | **1229** |
| T1 | - | 16 | 130 | 175 | 131 | 41 | 493 |
| T2-T4 | - | 69 | 182 | 141 | 228 | 116 | 736 |
| NA | 107 | 2 | 8 | 10 | 1 | 0 | 128 |
| **AGE**<br>**(Total)** | **107** | **87** | **320** | **326** | **360** | **157** | **1357** |
| < Median | 53 | 43 | 160 | 139 | 165 | 79 | 639 |
| >Median | 54 | 44 | 160 | 187 | 195 | 78 | 718 |

To determine whether the published signatures were suitable for systematic evaluation across the six TNBC cohorts, we next assessed signature quality as described in Methods section “Signature Scoring and Quality Control”^14^. All 62 signatures achieved goodness scores greater than 80% across the six cohorts, indicating that they were well represented, sufficiently variable, and internally coherent for robust downstream comparative analyses (Supplementary Figure S4E).

### Prognostic Signature Clusters Reveal Distinct Molecular and Biological Landscapes in TNBC

To identify biologically related subgroups among published TNBC prognostic signatures, we performed consensus k-means clustering of ssGSEA-based signature scores across the six discovery cohorts. This analysis identified three robust signature clusters characterized with distinct transcriptional programs (Figure 2A). Pathway enrichment analysis revealed marked biological differences between the clusters (Figure 2B). Cluster 1 was enriched for epithelial-mesenchymal transition (EMT), TNFα/NFκB signaling, IL6-JAK-STAT3 signaling, and p53 pathway activity, consistent with mesenchymal and inflammatory tumor states. Cluster 2 was characterized by proliferative programs, including E2F targets, G2M checkpoint, MYC targets, mTORC1 signaling, and glycolysis, reflecting highly proliferative basal-like tumors. In contrast, Cluster 3 exhibited enrichment for interferon-γ response, IL2-STAT5 signaling, KRAS signaling, and TGF-β pathways, indicative of immune-modulatory and stromal-associated transcriptional programs (Supplementary Table S1D). We next examined the distribution of these signature clusters across established TNBC molecular subtype classifications (Figure 2C). Cluster 1 was preferentially associated with LAR and mesenchymal-like tumors across PAM50^15,16^, TNBCtype-4^17,18^, and Burstein classification systems^19^, consistent with its EMT- and stromal-associated biology. Cluster 2 showed the strongest enrichment in basal-like tumors, particularly the BL1 subtype, reflecting the proliferative phenotype characteristic of basal TNBC^20^.

**Figure 2:**
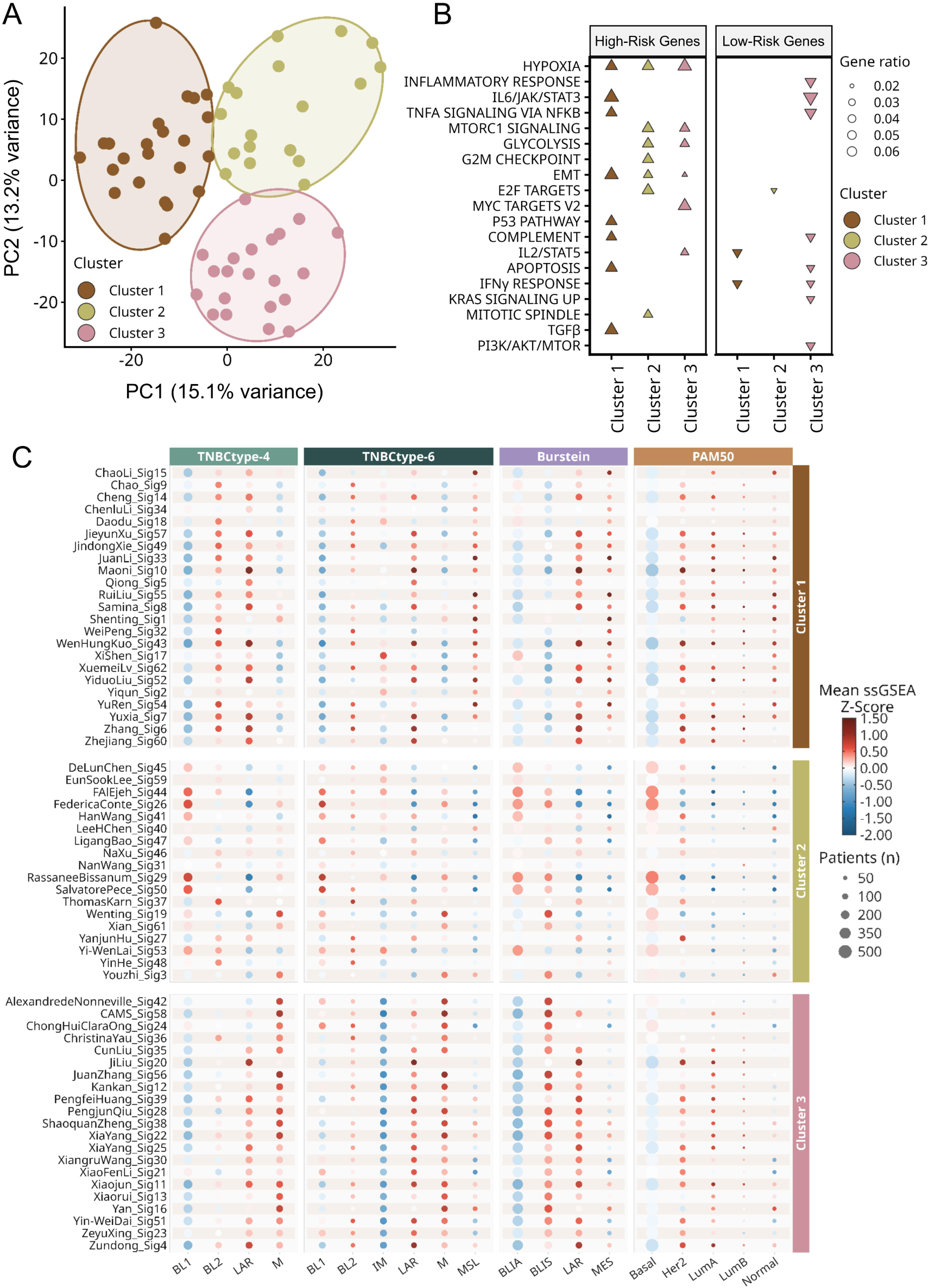
Consensus clustering of 62 prognostic signature scores reveals three distinct groups of signatures. A) Consensus K-means clustering (K = 3) of ssGSEA scores generated from 62 published TNBC prognostic signatures. B) Pathway enrichment analysis (Hallmark, MSigDB) of the union of high-risk and low-risk genes from prognostic signatures within each consensus cluster. High-risk and low-risk gene sets were analyzed separately. Dot size represents the gene ratio, and color denotes the consensus signature cluster. C) Mean ssGSEA Z-scores of signatures within each cluster across molecular subtypes defined by TNBCtype-4, TNBCtype-6, Burstein, and PAM50 classification systems. Dot size represents the number of patients within each subtype, and color indicates the mean ssGSEA Z-score.

Cluster 3 was enriched in immune-associated tumors and demonstrated overlap with specific LAR, mesenchymal, and BLIS subtype assignments, suggesting that immune activation represents a complementary biological axis that partially transcends conventional subtype boundaries.

Together, these findings demonstrate that the TNBC prognostic signature literature, despite substantial heterogeneity at the individual gene level, is organized around three biologically coherent programs: mesenchymal-inflammatory aggression (Cluster 1), proliferative-metabolic aggression (Cluster 2), and immune microenvironment activity (Cluster 3). This framework provided the rationale for subsequent meta-analytic analyses aimed at identifying signatures and genes that reproducibly capture prognostic information across these major biological themes.

### Cross-Cohort Meta-Analysis Reveals a Small Reproducibly Prognostic Subset Among 62 Published TNBC Signatures

A major challenge in evaluating transcriptomic prognostic signatures is the dependence of risk stratification on cohort-specific threshold selection. Across the six discovery cohorts (n = 1,357), the optimal threshold separating high- and low-risk patients varied substantially for many signatures despite identical scoring procedures, indicating marked inter-cohort heterogeneity in risk-score distributions (Figure 3; Supplementary Table S2A). To overcome this limitation, we developed a cross-threshold meta-analysis framework in which thresholds derived from each cohort were systematically applied across all six cohorts, and the resulting hazard ratios were integrated using random-effects meta-analysis (Refer to Methods Section: *Meta-Analysis Framework*). Using this framework, univariate analysis identified 17 of 62 signatures (27.4%) with reproducible prognostic associations for either EFS or OS (Supplementary Figure S5A-D; Supplementary Tables S2B-E). Of these, eight signatures (JiLiu_Sig20^21^, JuanZhang_Sig56^22^, ShaoquanZheng_Sig38^23^, XiangruWang_Sig30^24^, XiaYang_Sig22^25^, XiaYang_Sig25^26^, ZeyuXing_Sig23^27^, and Zundong_Sig4^28^) remained consistently significant across all four scoring-endpoint combinations, demonstrating that only a small subset of published signatures exhibits robust prognostic behavior independent of scoring strategy.

**Figure 3:**
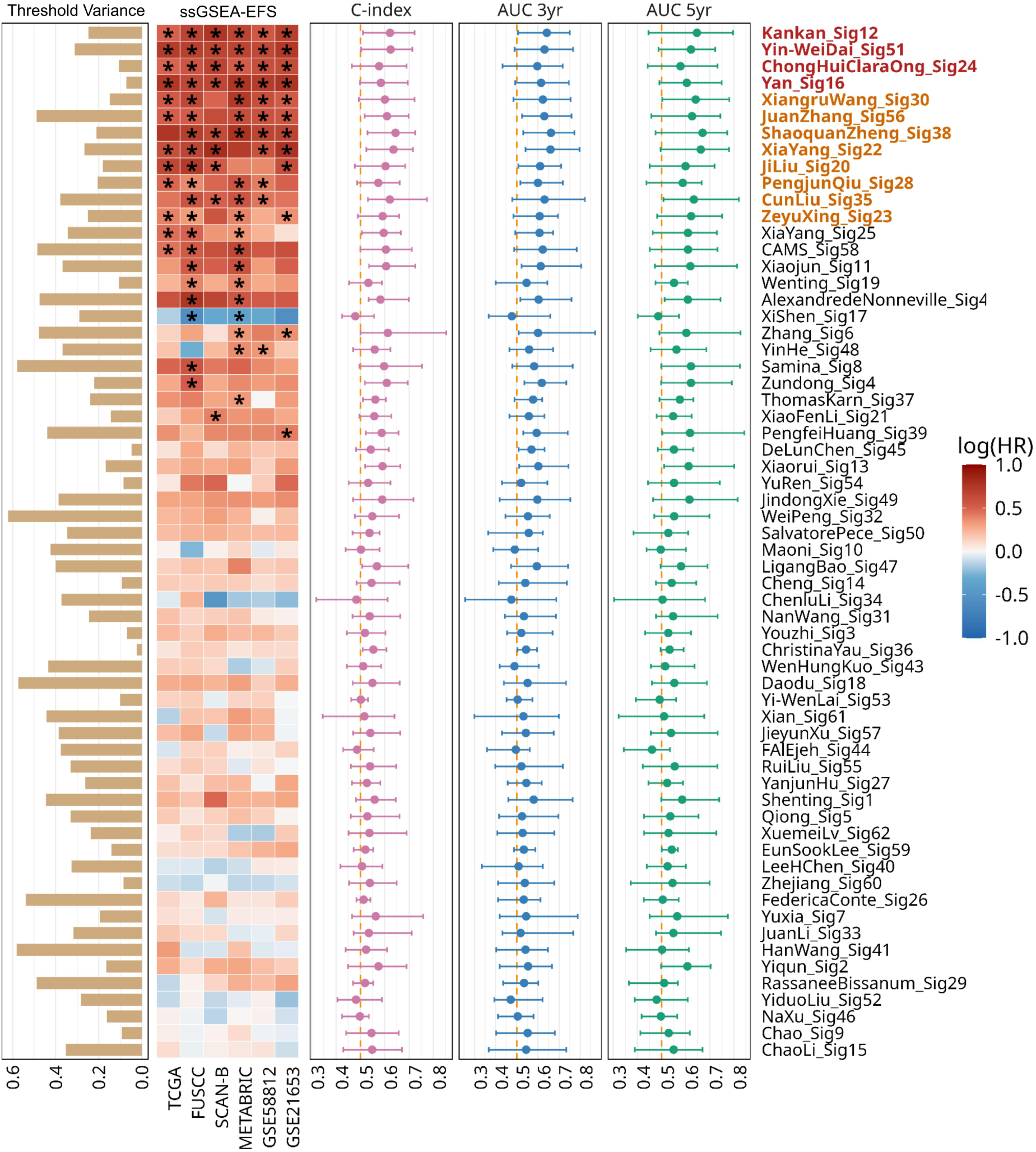
Multivariable Random Effects Meta-Analysis of published prognostic signatures for EFS. Variability in median cohort thresholds (ssGSEA-derived) of six discovery cohorts. (Refer to Methods Section: Meta-Analysis Framework). The heatmap shows pooled hazard ratios (HRs) for each signature (rows) and threshold set (columns); color indicates log(HR), and asterisks denote significant meta-analysis results (FDR < 0.05). Forest plots present pooled C-index, 3-year AUC, and 5-year AUC with 95% confidence intervals. Signature names in red were significant across all cohorts, while coral indicates significance in at least four cohorts.

Adjustment for age, tumor size, and lymph node status substantially reduced the number of reproducible signatures. Only 4 of 62 signatures (6.5%) remained consistently significant across all threshold schemes for EFS: Kankan_Sig12^29^ (HR range 1.64-2.14), Yin-WeiDai_Sig51^30^ (HR range 1.92-2.10), ChongHuiClaraOng_Sig24^31^ (HR range 1.63-1.98), and Yan_Sig16^32^ (HR range 1.93-2.16) (Figure 3; Supplementary Table S2F). An additional eight signatures retained significance in at least four of six threshold schemes (highlighted in coral in Figure 3). Nevertheless, even these reproducible signatures demonstrated only modest discriminatory performance, with mean C-indices ranging from 0.54 to 0.62, indicating that statistical reproducibility alone does not ensure clinically useful prediction.

The limited reproducibility of published signatures was consistent across alternative analytical frameworks. Multivariable significance was retained by only 5 of 62 signatures for SingScore-EFS, 2 of 62 for ssGSEA-OS, and 2 of 62 for SingScore-OS (Supplementary Figure S6A-C; Supplementary Tables S2G-I). Similarly, among the 38 signatures with published coefficient-based formulas, weighted risk-score analysis identified only 10 signatures significant for EFS and one signature for OS after multivariable adjustment (Supplementary Figure S7A-B; Supplementary Tables S2J-K), with limited overlap with the unweighted analyses. Direct comparison across scoring methods further emphasized the scarcity of broadly reproducible signatures. Within the EFS analyses, only Kankan_Sig12^23^, Yan_Sig16^29^, and Yin-WeiDai_Sig51^30^ remained significant under both ssGSEA and SingScore scoring, whereas ShaoquanZheng_Sig38^32^ was the only reproducible signature for OS. Importantly, no published signature remained consistently significant across all four scoring-endpoint combinations (Supplementary Figure S6D). Together, these findings demonstrate that the prognostic performance of existing TNBC signatures depends strongly on analytical choices, including threshold selection, scoring strategy, and clinical endpoint, motivating the development of a new multi-cohort-derived prognostic classifier. Because rankings based on hazard ratio, C-index, and AUROC were highly concordant, subsequent analyses focus on hazard ratio-based benchmarking for clarity.

### Derivation of MetaSig: A Multi-Cohort Prognostic Classifier for TNBC

To overcome the limited cross-cohort reproducibility of existing prognostic signatures, we developed a multi-cohort gene discovery framework that retained only genes demonstrating statistically significant survival associations with concordant hazard ratio directions across all six discovery cohorts. To minimize bias arising from differences in cohort size and event rate, we identified a Directionally Concordant Gene set (DCGS) comprising genes that demonstrated consistent hazard ratio direction across all six discovery cohorts. To account for dataset-specific expression thresholds, gene selection was performed using a resampling-based strategy, retaining only genes with reproducible prognostic effects across cohorts (see Methods: Multi-Cohort Gene Discovery and Model Training).

This framework identified 15 directionally concordant gene set associated with EFS (DCGS-EFS): *ALDOA, FSTL3, INTS11, PPP1R13B, PPP1R13L, SPINT1, STC1, COL6A6, DEF6, GBP5, ICOS, NFE2L3, PVRIG, TNFSF13B,* and *TTC39C* (Figure 4A). These genes segregated into two opposing biological programs. Poor-prognosis genes were broadly associated with metabolic adaptation, tumor progression, and stromal remodeling: *ALDOA* is a key regulator of glycolytic adaptation under hypoxia^33^; *PPP1R13L (IASPP)* suppresses p53-dependent transcription and has been linked to therapy resistance^34^; and *SPINT1* has been implicated in tumor invasion and metastatic progression^35^. Consistent with this, *FSTL3* and *STC1* are associated with stromal remodeling, immune exclusion, angiogenesis, and resistance to apoptosis^36,37^. By contrast, favorable prognostic genes reflected coordinated adaptive immune responses: I*COS, GBP5, DEF6*, and *TNFSF13B* regulate T-cell activation, interferon signaling, inflammasome function, and B-cell survival within the tumor microenvironment^38–41^. Notably, these functional annotations closely paralleled the compartment-specific expression patterns observed in the single-cell analyses described below. A parallel analysis of overall survival identified 20 directionally concordant gene set (DCGS-OS), including *CD274, COL6A6, CYP27A1, EGFL6, FAM117A, IL27RA, LPIN2, MICB, NFE2L3, PVRIG, RUNX3, SOCS1, HIP1R, LYPLA2, SLC52A3, SLC6A1, STC1, SULT2B1, TRIP10*, and *ULK1* (Supplementary Figure S8A; Supplementary Table S3A).

**Figure 4:**
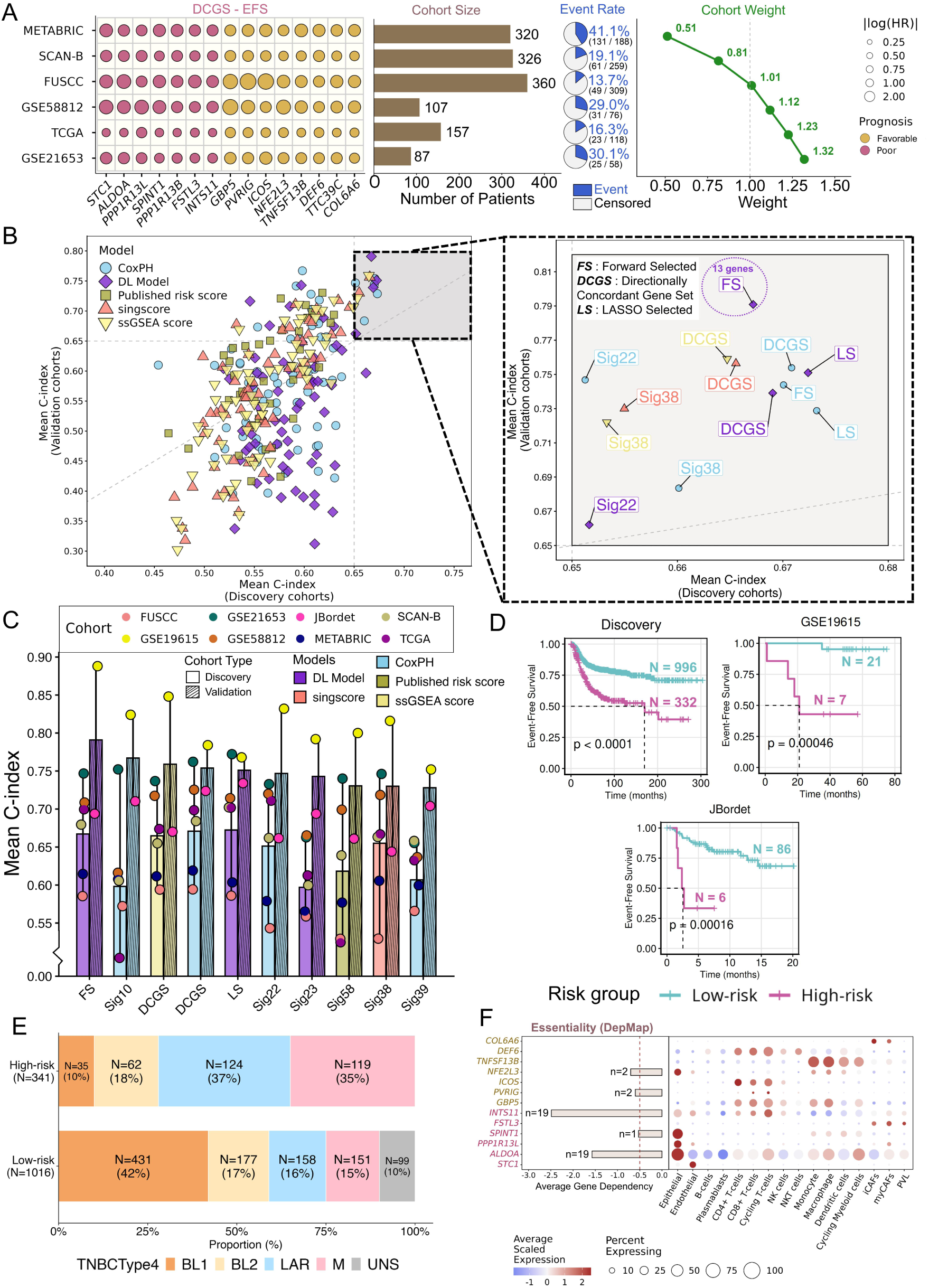
Development and validation of MetaSig, a multi-cohort prognostic classifier for TNBC. A) Identification of the Directionally Concordant Gene Set (DCGS-EFS) across the six discovery cohorts and the cohort-balanced weighting scheme used for model training. Dot size represents the pooled hazard ratio (HR) magnitude and color indicates prognostic direction. The accompanying panel summarizes cohort characteristics, including sample size, event rate, and the relative cohort weights assigned during model development to balance differences in cohort size and outcome frequency. B) Performance of prognostic modeling strategies comparing the 62 published prognostic signatures, the Directionally Concordant Gene Sets (DCGS), and their Forward-Selected (FS) and LASSO-Selected (LS) reduced models using Cox proportional hazards (CoxPH) and DeepSurv. Mean concordance index (C-index) in the six discovery cohorts is plotted against the mean C-index in the external validation cohorts. The inset highlights the highest-performing models. MetaSig-EFS, derived from the FS-DCGS-EFS gene set and trained using DeepSurv, demonstrated the best overall predictive performance. C) Mean C-index of the top-performing models across individual discovery and validation cohorts. Bars indicate discovery performance, striped bars indicate validation performance, and points represent cohort-specific C-indices. D) Kaplan-Meier curves showing event-free survival stratified by MetaSig-EFS risk groups in the combined discovery cohort and external validation cohorts. Risk groups were defined using the discovery cohort threshold (top 25% high-risk) and applied unchanged to validation cohorts. E) Distribution of TNBCtype-4 molecular subtypes across MetaSig-EFS low- and high-risk groups. F) Mean gene dependency scores from DepMap and single-cell expression of MetaSig-EFS genes across cell types in the TNBC tumor microenvironment. Genes are colored according to their prognostic direction in the DCG-EFS signature (Figure 4A), with red indicating poor-prognosis genes and gold representing favorable-prognosis genes.

Four of these genes - *COL6A6, NFE2L3, PVRIG, and STC1* - were shared between the EFS and OS signatures, suggesting conserved prognostic relevance across both clinical endpoints. LASSO and forward stepwise selection were then applied to derive reduced EFS and OS feature sets for model development (Supplementary Table S3B).

With these candidate gene sets in hand, we next asked which modeling strategy would best translate them into a clinically useful score. We evaluated Cox proportional hazards^42^ and weighted DeepSurv^43^ models using all 62 published TNBC prognostic signatures, as well as the 15-gene DCGS-EFS and 20-gene DCGS-OS signatures and their respective forward-selected (FS) and LASSO-selected (LS) derivatives. Model training incorporated a cohort-balanced weighting scheme together with a leave-one-cohort-out (LOCO) optimization framework to maximize generalizability while preventing larger or event-rich cohorts from disproportionately influencing model development (see Methods: Cohort-Balanced Model Training; Supplementary Table S3C). Model performance was benchmarked against ssGSEA, SingScore, and the original coefficient-based risk scores where available, yielding 294 models/scoring strategies for each survival endpoint; discovery cohort characteristics are summarized in Figure 4A and Supplementary Figure S8A.

Under the leave-one-cohort-out (LOCO) optimization framework (see Methods: Cohort-Balanced Model Training), a forward-selected 13-gene DeepSurv model derived from the 15-gene DCGS-EFS signature achieved the best overall performance for EFS (Figure 4B-C), whereas a LASSO-selected 7-gene Cox proportional hazards model derived from the 20-gene DCGS-OS signature performed best for OS (Supplementary Figure S8B-C; full metrics in Supplementary Tables S3D-E). These optimized signatures were designated MetaSig-EFS and MetaSig-OS, respectively.

Across the discovery cohorts, MetaSig-EFS achieved a mean LOCO C-index of 0.67 (range: 0.59-0.74) (Figure 4C; Supplementary Table S3D). In external validation, it achieved concordance indices of 0.89 in GSE19615 and 0.69 in JBordet, with corresponding 3- and 5-year time-dependent AUROCs of 0.89 and 0.96 in GSE19615 and 0.80 and 0.71 in JBordet, respectively. MetaSig-OS performed comparably, achieving a mean LOCO C-index of 0.66 (range: 0.601-0.761) across the discovery cohorts and validation concordance indices of 0.71 in JBordet, with corresponding 3-and 5-year AUROCs of 0.90 and 0.76 (Supplementary Figure S8C; Supplementary Table S3E).

To translate these continuous scores into a clinically actionable tool, we derived fixed risk-score thresholds from the discovery cohorts using DeepSurv outputs, classifying patients in the highest quartile as high risk and applying these thresholds unchanged to the validation cohorts. The MetaSig-EFS threshold (0.06) reproducibly separated patients into distinct risk groups in both external cohorts (log-rank P = 0.00046, GSE19615; P = 0.00016, JBordet), and the MetaSig-OS threshold (1.31) did the same in the JBordet validation cohort (log-rank P < 0.001) (Figure 4D; Supplementary Figure S8D; Supplementary Table S3F). Together, these results show that MetaSig functions both as a continuous prognostic model and as a robust binary risk-classification tool.

Having established its prognostic performance, we next asked what biology underlies MetaSig’s risk stratification. Pathway enrichment revealed a clear dichotomy between risk groups: high-risk tumors were enriched for epithelial-mesenchymal transition, hypoxia, glycolysis, androgen signaling, and lipid metabolism, while low-risk tumors preferentially activated interferon responses, inflammatory signaling, complement activation, and cytokine-mediated immune pathways (Supplementary Table S3H-I) - indicating that MetaSig captures a transition from immune-active to metabolically aggressive tumor states.

MetaSig also added prognostic resolution beyond established TNBC molecular subtypes. High-risk tumors were significantly enriched within the luminal androgen receptor (LAR) and mesenchymal subtypes, while low-risk tumors were predominantly basal-like (chi-square test, *p* < 2.2 × 10⁻¹⁶; Cramér’s *V* = 0.396 for MetaSig-EFS and *p* < 2.2 × 10⁻¹⁶; Cramér’s *V* = 0.298 for MetaSig-OS) (Figure 4E; Supplementary Figure S8E). Even so, both favorable- and poor-prognosis tumors occurred within every subtype, and MetaSig significantly stratified survival across nearly all Burstein, TNBCtype-4, and TNBCtype-6 subtype groups (Supplementary Figure S9-S10; Supplementary Table S3G).

To understand what drives this subtype-independent stratification, we examined the biological basis of risk differences within each TNBCtype-4 subtype. Across BL1, BL2, LAR, and M tumors, high-risk groups consistently showed enrichment of metabolic and stress-associated programs - androgen response, fatty acid metabolism, cholesterol homeostasis, glycolysis, oxidative phosphorylation, and hypoxia (Supplementary Figure S11A-B) - whereas low-risk tumors were characterized by immune and inflammatory programs, including interferon-α and interferon-γ responses, IL6/JAK/STAT3 signaling, TNFα signaling via NF-κB, inflammatory response, and complement activation (Supplementary Table S3H-I). These patterns held across both MetaSig-EFS and MetaSig-OS and were most pronounced in the LAR subtype, where high-risk tumors showed strong activation of androgen response, fatty acid metabolism, and cholesterol homeostasis programmes.

To resolve which cell populations underlie these transcriptomic programs, we analyzed a single-cell RNA-seq dataset of 25,972 cells from seven treatment-naïve primary TNBC tumors^44^, resolving 16 epithelial, immune, stromal, and vascular cell populations after quality control and batch correction (Supplementary Figure S13A-D). MetaSig-EFS genes showed striking compartment-specific expression across the tumor microenvironment: favorable prognostic genes (*ICOS, GBP5, DEF6, PVRIG*, and *TNFSF13B*) were predominantly expressed in T-cell and other immune populations, while adverse prognostic genes localized to distinct tumor-associated compartments - *ALDOA*, *PPP1R13L*, and *SPINT1* in malignant epithelial cells, *FSTL3* in cancer-associated fibroblasts (CAFs) and perivascular-like (PVL) cells, and *STC1* in endothelial cells (Figure 4F). MetaSig-OS genes followed a comparable pattern, with favorable prognostic genes (*LPIN2*, *SOCS1*, and *NFE2L3*) enriched in immune populations and adverse genes localizing to epithelial, endothelial, and B-cell compartments (Supplementary Figure S8G).

Building on this compartment mapping, we then asked whether these genes were also functionally required by tumor cells, using CRISPR-based dependency profiles across 19 TNBC cell lines. Five MetaSig-EFS genes (*ALDOA, NFE2L3, SPINT1, INTS11,* and *PVRIG*) showed dependency scores consistent with selective essentiality in at least one TNBC cell line, while *NFE2L3* was the only MetaSig-OS gene meeting this criterion (Figure 4F; Supplementary Figure S8F). Together, these findings show that MetaSig genes represent biologically interpretable, compartment-specific transcriptional programs with functional relevance in TNBC.

Finally, we tested whether MetaSig’s prognostic value held independently of standard clinicopathological variables. For MetaSig-EFS, multivariable Cox analysis confirmed the signature as an independent predictor of outcome (HR 2.03, 95% CI 1.58-2.59, P < 0.001), alongside lymph node status (HR 2.24, P < 0.001) and tumor size (HR 1.41, P = 0.009); age was not independently associated with EFS (Supplementary Figure S12A). Similar results were observed for MetaSig-OS, though age remained independently associated with overall survival (Supplementary Figure S12E). MetaSig also consistently stratified survival within each clinical subgroup defined by lymph node status, tumor size, and age (Supplementary Figure S12B-D, F-H), confirming that its prognostic value is independent of established clinicopathological variables.

### Master Regulator Analysis Reveals Distinct Regulatory Programs Underlying MetaSig Risk States

To identify the transcriptional programs underlying the MetaSig risk states, we inferred master regulator activity using VIPER. We identified 54 significant master regulators for MetaSig-EFS and 53 for MetaSig-OS (FDR <0.05), of which 41 were shared between both prognostic frameworks, indicating that recurrence and overall survival are governed by highly conserved regulatory programs.

Several regulators, including *PRDM1, FLI1, IRF1, EOMES and STAT4*, demonstrated reproducible activity across external validation cohorts, supporting the robustness of the inferred regulatory architecture (Supplementary Table S3J). The regulatory landscape segregated into two distinct transcriptional states associated with patient outcome (Figure 5A).

**Figure 5:**
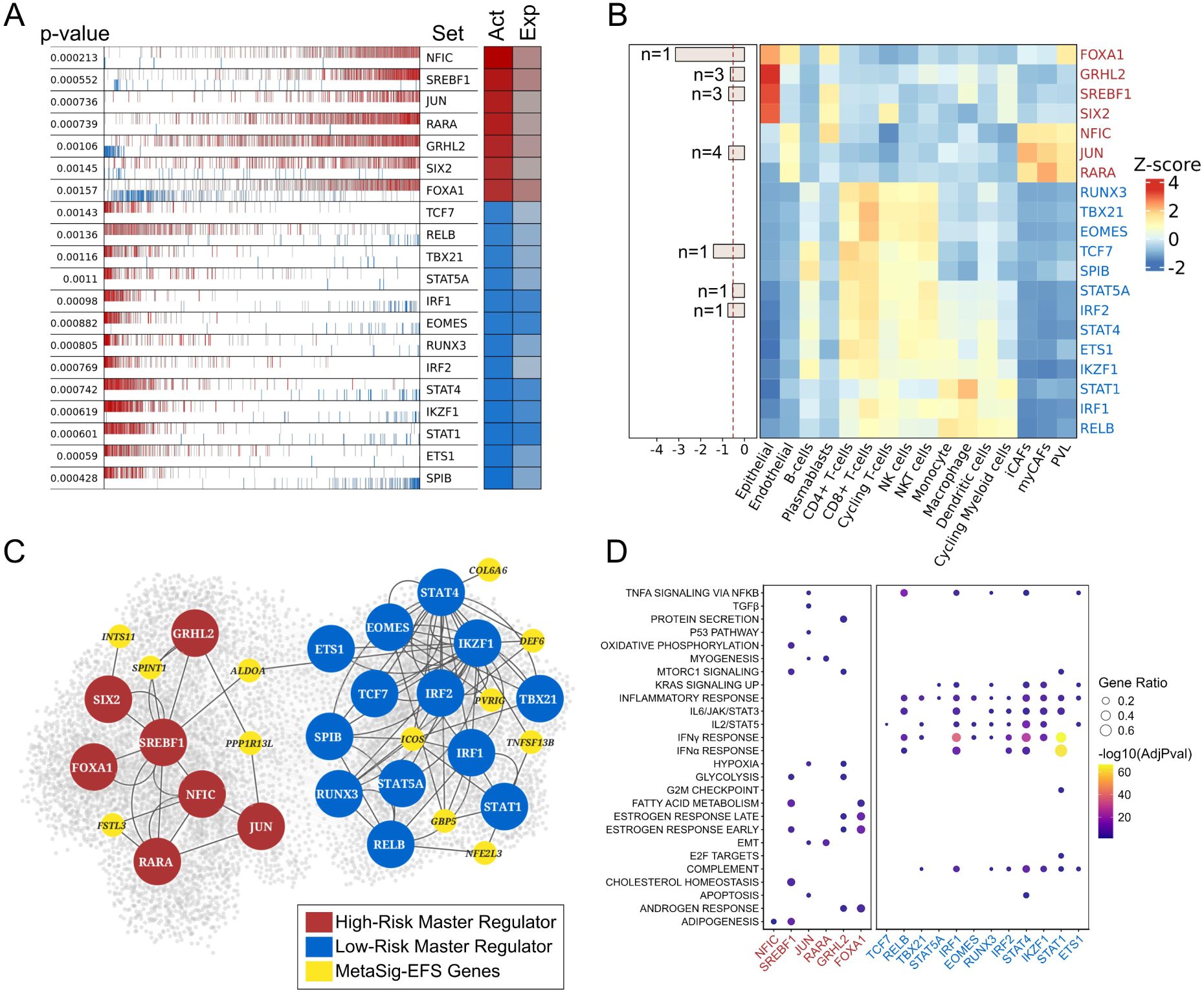
Upstream Regulatory Architecture of MetaSig-EFS. A) Plot showing the top 20 activated and repressed master regulators (MRs) identified between high- and low-risk patients using VIPER analysis. The leftmost column indicates the significance (p-value) of each regulon. The central barcode-like panel represents the distribution of activated (red) and repressed (blue) target genes ranked according to the differential expression signature (left: most down-regulated; right: most up-regulated). The “Act” column represents inferred regulon activity, while the “Exp” column indicates the expression level of the corresponding transcription factor. B) Heatmap showing the mean scaled AUCell regulon activity scores of the MetaSig-EFS master regulators across cell types in the TNBC single-cell dataset. Colors represent the mean scaled AUCell score for each regulon within each cell type. The horizontal bar plot summarizes gene essentiality based on DepMap dependency scores across 19 TNBC cell lines. Bar length corresponds to the mean dependency score, while *n* indicates the number of TNBC cell lines in which the corresponding master regulator was classified as essential. C) Network representation of MRs and their association with MetaSig-EFS genes. Red nodes indicate MRs activated in high-risk patients, whereas blue nodes represent MRs activated in low-risk patients. Yellow nodes denote MetaSig-EFS genes connected to the corresponding regulatory network. Edges represent inferred regulatory interactions between transcription factors and target genes. D) Dot plot depicting pathways represented by high- and low-risk patient-specific MRs. Regulons activated in low-risk patients were predominantly enriched for immune-related and inflammatory pathways, whereas high-risk regulons showed enrichment for metabolic, proliferative, and EMT-associated programs. Dot size represents gene ratio, and color intensity indicates significance (-log10 adjusted p-value).

High-risk tumors were characterized by activation of *NFIC, SREBF1, JUN, RARA, GRHL2, SIX2, and FOXA1*, whereas low-risk tumors exhibited increased activity of *TCF7, RELB, TBX21, STAT5A, IRF1, EOMES, RUNX3, IRF2, STAT4, IKZF1, STAT1, ETS1*, and *SPIB*. A similar pattern was observed for MetaSig-OS, highlighting a high degree of concordance between the regulatory programs associated with recurrence and survival (Supplementary Figure S14A).

To determine the cellular context of these regulatory programs, we mapped MR expression onto the single-cell atlas (Figure 5B; Supplementary Figure S14B). High-risk regulators were predominantly expressed in malignant epithelial cells and, for several regulators including NFIC, JUN, and RARA, extended into cancer-associated fibroblast (CAF) and perivascular-like (PVL) compartments. In contrast, low-risk regulators were enriched across immune populations, including T cells, NK cells, monocytes, and macrophages, consistent with the immune-associated phenotype identified by MetaSig. Several regulators, including FOXA1, GRHL2, SREBF1, JUN, TCF7, STAT5A, and IRF2, also exhibited dependency scores consistent with selective essentiality in TNBC cell lines, suggesting that these regulatory programs contribute to tumor cell fitness in specific TNBC contexts (Figure 5B).

We next investigated how these master regulators coordinate expression of MetaSig genes. Integration of master regulators with MetaSig-EFS genes revealed a highly organized regulatory network connecting prognostic genes to distinct biological processes (Figure 5C). Favorable prognostic genes, including ICOS, GBP5, DEF6, and TNFSF13B, were embedded within immune-associated regulatory modules centered on STAT1, STAT4, IRF1, and IKZF1. In contrast, adverse prognostic genes such as ALDOA, PPP1R13L, and SPINT1 were linked to regulators governing metabolic adaptation, epithelial-mesenchymal transition, hypoxia, and hormone-responsive signaling (Figure 5D). Notably, ALDOA occupied a central network position connecting immune- and tumor-associated regulatory modules, suggesting a potential interface between metabolic reprogramming and immune regulation.

Consistent with these network-level observations, low-risk regulators were strongly enriched for interferon signaling, inflammatory response, IL2-STAT5 signaling, IL6-JAK-STAT3 signaling, and TNFα signaling via NFκB, whereas high-risk regulators were associated with cholesterol homeostasis, fatty acid metabolism, glycolysis, oxidative phosphorylation, mTORC1 signaling, androgen response, and epithelial-mesenchymal transition. Similar pathway enrichment was observed for MetaSig-OS (Supplementary Figure S14C-D; Supplementary Table S3K), indicating that both prognostic models-converge on a common regulatory architecture linking immune activation with favorable outcome and metabolic reprogramming with aggressive disease.

### Single-Cell Dissection of TNBC Prognostic Risk Reveals High-Risk Epithelial States and Actionable Therapeutic Vulnerabilities

Further, to uncover the cellular correlates of prognostic heterogeneity, we used a phenotype-guided bulk-to-single-cell regression approach^45^ to map bulk-derived high-and low-risk phenotypes onto the single-cell transcriptomic landscape, thereby identifying the cellular populations underlying each prognostic state. This analysis identified 2,033 high-risk (7.83% of all cells) and 2,465 low-risk (9.49% of all cells) cells for MetaSig-EFS. For MetaSig-OS, 1,977 high-risk (7.61%) and 2,293 low-risk (8.83%) cells were identified, while the remaining cells were classified as background by the Scissor algorithm (Supplementary Table S3M). Scissor reliability testing confirmed the robustness of the identified prognostic cell populations (reliability statistic = 0.94, *P* < 0.05) The resulting cellular distributions revealed a striking segregation of risk-associated cells across the tumor ecosystem (Figure 6A-B; Supplementary Figure S17A, B).

**Figure 6:**
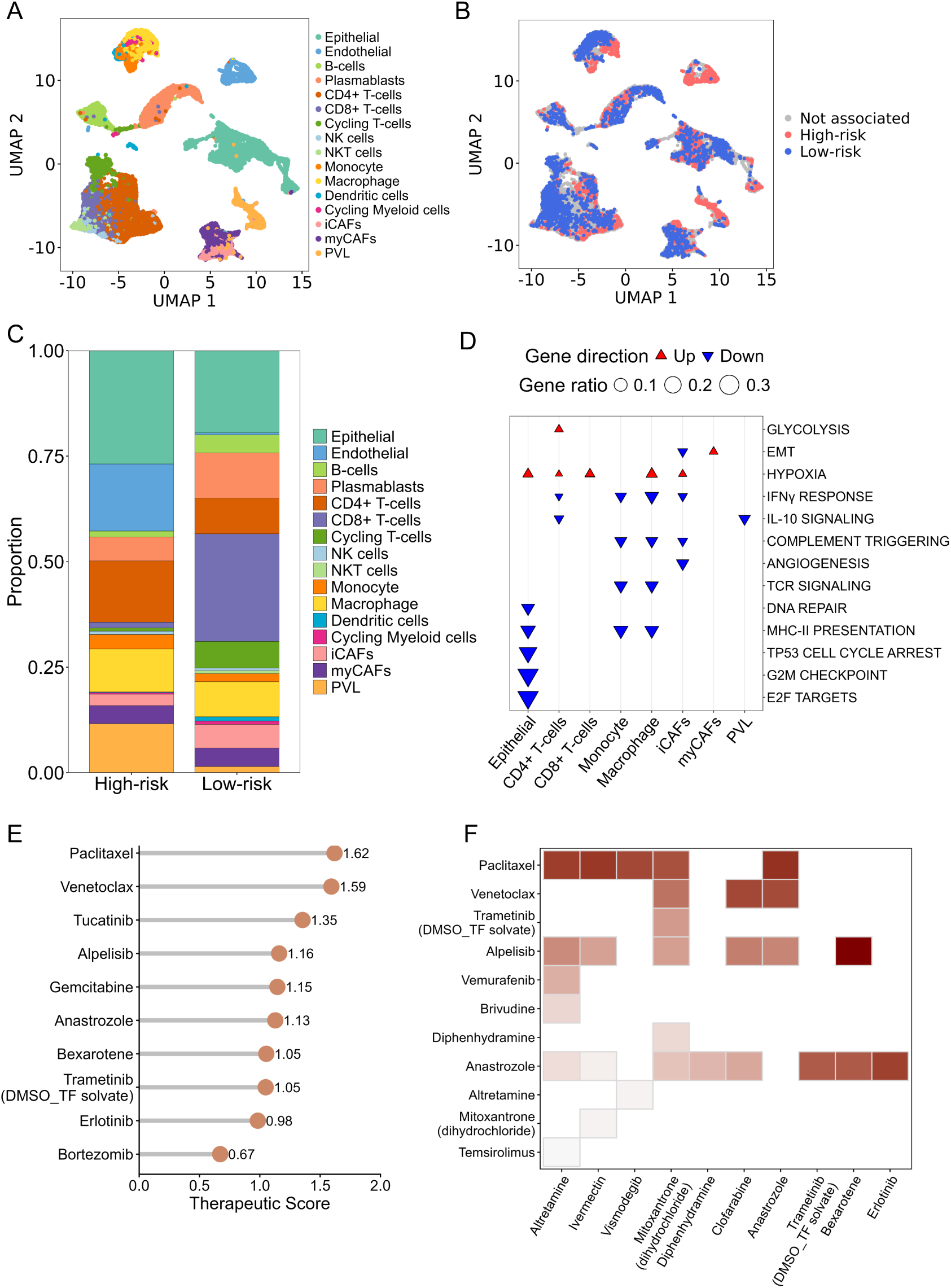
Risk-Associated Cell States and Candidate Therapeutics for MetaSig-EFS. **(A)** UMAP visualization showing the distribution of annotated cell types in the TNBC single-cell dataset. B) UMAP depicting the distribution of cells associated with high-risk and low-risk patients across the single-cell landscape. Red cells represent populations associated with high-risk patients, blue cells represent populations associated with low-risk patients, and grey cells indicate populations not significantly associated with either risk group. C) Stacked bar plot showing the proportional distribution of annotated cell types among cells associated with high-risk and low-risk patients. D) Pathway analysis comparing high-risk and low-risk associated cells across individual cell types. Red upward triangles indicate pathways significantly up-regulated in high-risk cells, whereas blue downward triangles represent pathways enriched in low-risk cells. Dot size corresponds to gene ratio, reflecting the proportion of pathway-associated genes contributing to the signal. E) Top FDA-approved therapeutic drugs identified through drug repurposing analysis using differential gene expression signatures derived from epithelial cells associated with high- versus low-risk patients in the Tahoe perturbation dataset. F) Heatmap showing significant candidate drug combinations predicted for high-risk patients. Colored intensity represents the combination therapeutic score, with darker shades indicating stronger predicted synergistic therapeutic potential.

High-risk cells were predominantly localized within malignant epithelial and endothelial compartments, whereas low-risk cells were enriched in adaptive immune populations, particularly T and B cells (Figure 6C; Supplementary Figure S15C). These differences were highly significant (chi-square test, *p* < 2.2 × 10⁻¹⁶), with large effect sizes for both MetaSig-EFS (Cramér’s *V* = 0.53) and MetaSig-OS (Cramér’s *V* = 0.54), demonstrating a strong association between prognostic risk and the cellular composition of the tumor microenvironment. Notably, highly similar cellular landscapes were identified for both MetaSig-EFS and MetaSig-OS despite being derived from independent clinical endpoints, with high-risk states consistently enriched for epithelial, endothelial and stromal compartments and low-risk states dominated by adaptive immune populations. While bulk transcriptomic analyses defined the prognostic programs associated with MetaSig risk, bulk-to-single-cell integration resolved the cellular basis of these programs, revealing that adverse prognostic states arise predominantly from tumor- and stromal-associated compartments, whereas favorable-risk states are primarily associated with immune cell populations.

The biological pathways represented by these cellular states further reinforced their distinct prognostic roles. High-risk cells exhibited enrichment of glycolysis and hypoxia, whereas low-risk cells were characterized by activation of interferon signaling, antigen presentation, complement pathways, and broader immune response programs (Figure 6D; Supplementary Table S3M). Comparable pathway enrichments were observed in the MetaSig-OS analysis, indicating convergence on a common set of biological processes despite differences in clinical endpoints (Supplementary Figure S15D; Supplementary Table S3N).

The biological pathways represented by these cellular states further reinforced their distinct prognostic roles. High-risk cells exhibited enrichment of glycolysis and hypoxia, whereas low-risk cells were characterized by activation of interferon signaling, antigen presentation, complement pathways, and broader immune response programs (Figure 6D; Supplementary Table S3M). Comparable pathway enrichments were observed in the MetaSig-OS analysis, indicating convergence on a common set of biological processes despite differences in clinical endpoints (Supplementary Figure S15D; Supplementary Table S3N).

Collectively, these findings demonstrate that the bulk-derived prognostic signatures capture fundamentally different tumor ecosystem states. Poor prognosis is associated with epithelial-dominant, metabolically active tumor programs, whereas favorable outcomes are characterized by immune-enriched micro-environments marked by robust inflammatory and antigen-presentation responses. These observations provide a cellular framework linking transcriptomic risk stratification to the biological organization of the TNBC tumor microenvironment.

Next, we sought to identify compounds capable of reversing the high-risk transcriptional program of epithelial cells, the predominant population within the high-risk compartment. Using disease-specific signatures, we performed a computational drug repurposing analysis with the Tahoe-100M single-cell perturbation atlas^46^ and identified FDA-approved drugs predicted to reverse the high-risk epithelial transcriptional state (Supplementary Table S3O). Among the top-ranked compounds identified for MetaSig-EFS (Figure 6E; Supplementary Table S3P), Paclitaxel achieved the highest reversal score (1.62), followed by Venetoclax (1.59). The prioritization of Paclitaxel, a standard first-line chemotherapeutic agent for TNBC, provides an internal positive control supporting the biological validity of the transcriptomic reversal framework. Although *BCL2* expression was low in the high-risk epithelial population, Venetoclax has been reported to exert *BCL2*-independent effects on mitochondrial metabolism and the integrated stress response, providing a potential explanation for its prioritization^47^.

Tucatinib (1.35), a HER2-selective tyrosine kinase inhibitor^48^, ranked third. Although TNBC is traditionally classified as HER2-negative, recent studies have identified a molecularly distinct HER2-low/LAR subgroup that may benefit from HER2-targeted therapies^49^. Additional highly ranked candidates included Alpelisib (1.16), targeting the PI3K pathway^50^, and Gemcitabine (1.15), a nucleoside analogue with established activity in metastatic TNBC^51^. Several compounds, including Paclitaxel, Venetoclax, Tucatinib, Alpelisib and Gemcitabine, were also prioritized in the MetaSig-OS analysis (Supplementary Figure S15E; Supplementary Table S3P), indicating that both prognostic models converge on common therapeutic vulnerabilities in high-risk TNBC epithelial cells.

Beyond single-agent prioritization, we evaluated pairwise drug combinations among the top-ranked candidates to identify combinations capable of achieving enhanced reversal of the high-risk disease program. This analysis identified several high-scoring drug pairs involving Paclitaxel, Venetoclax and Anastrozole demonstrating the strongest therapeutic potential (Figure 6F; Supplementary Table S3Q). These findings suggest that simultaneous targeting of distinct oncogenic and survival-associated transcriptional pathways may provide greater efficacy in reversing the high-risk epithelial state than single-agent treatment alone. In the MetaSig-OS analysis, several overlapping therapeutic candidates were identified, including Paclitaxel and Venetoclax, alongside additional candidates such as Trametinib and Crizotinib, which emerged both as high-ranking single agents and combination partners, further supporting convergence on shared oncogenic signaling and survival dependencies in high-risk TNBC (Supplementary Figure S15F; Supplementary Table S3Q).

Consistent with the biological relevance of the framework, established breast cancer therapeutics were preferentially ranked among the top candidates, with 8/12 and 6/12 positive-control drugs falling within the top 50% of MetaSig-EFS and MetaSig-OS predictions, respectively (Supplementary Figure S16A-B). Collectively, the single-agent and combination analyses converged on a mechanistically diverse set of candidate therapeutics spanning cytotoxic chemotherapy, kinase inhibition, apoptotic modulation, and transcriptional reprogramming. These compounds represent potential candidates for downstream experimental validation in HS-578T cell line and additional TNBC models to evaluate their ability to reverse the transcriptional features associated with the high-risk disease state.

### Risk Prediction server

To facilitate reproducibility and independent application of the proposed models, we developed a web-based application for calculating risk scores based on the developed prognostic gene signatures. The tool was implemented using the Flask (v3.1.3) framework in Python (v3.11.9) and enables users to upload a gene expression matrix (FPKM or TPM). If the uploaded values are not log_2_ (*x* + 1)-normalized, the application automatically performs log_2_ (*x* + 1) transformation prior to analysis. The processed expression data are then used to compute individualized risk scores and assign patients to the corresponding risk groups based on predefined thresholds. The application is publicly accessible at https://metasig.igib.res.in.

## Discussion

Despite the large number of transcriptomic prognostic signatures proposed for triple-negative breast cancer (TNBC), none has been incorporated into routine clinical practice^52^. Systematic evaluation of 62 published TNBC prognostic signatures revealed limited reproducibility across independent cohorts despite their reported prognostic performance in individual studies. Yet this apparent inconsistency was accompanied by striking biological convergence. Prognostic gene signatures with minimal overlap in gene composition repeatedly captured the same immune, proliferative and stromal programs. This indicates that the major obstacle to clinical translation has not been the absence of reproducible biology, but the difficulty of identifying molecular surrogates that remain stable across diverse patient populations.

The limited reproducibility of published prognostic gene signatures appears to arise, at least in part, from their analytical design. Many published models were developed using overlapping discovery cohorts particularly TCGA^6^ and METABRIC^7^, optimized with cohort-specific coefficients and evaluated using a single scoring strategy. Under these conditions, even relatively modest differences in patient composition, expression distributions or threshold selection can substantially influence model performance. This likely explains why signatures that appeared highly prognostic in their original reports frequently showed reduced performance when evaluated under standardized conditions. Importantly, adjustment for tumor size, lymph node status and age further reduced the number of independently prognostic models, indicating that much of the predictive information captured by existing prognostic gene signatures overlaps with established clinicopathological variables. These findings argue that reproducibility across independent populations should become the primary benchmark for prognostic biomarker development, rather than performance within individual datasets.

Perhaps the most interesting observation was the disconnect between gene composition and biological function. Individual prognostic gene signatures shared remarkably few genes, yet repeatedly converged on the same transcriptional programs. Mesenchymal remodeling, metabolic adaptation, cell-cycle progression and adaptive immune activation emerged consistently irrespective of the genes comprising each signature. This suggests that prognosis in TNBC is encoded principally at the level of biological networks rather than individual transcripts^53^. Different studies therefore appear to identify alternative molecular representations of the same underlying tumor states. This observation also provides a plausible explanation for the long-standing paradox that signatures with minimal gene overlap often achieve similar prognostic performance. From a biomarker perspective, the challenge is therefore less about identifying new prognostic genes than about recognizing the conserved biological processes that they collectively represent.

These findings provided the rationale for developing MetaSig. Rather than refining existing prognostic gene signatures, we performed transcriptome-wide survival analyses independently within each discovery cohort and retained only genes showing significant and concordant associations with outcome across all six datasets. This stringent strategy minimized cohort-specific effects while prioritizing reproducible prognostic signals. MetaSig was derived independently of the published prognostic gene signatures evaluated here, yet recapitulated the same immune, proliferative and stromal programs identified during benchmarking.

We next evaluated whether these reproducible features could be more effectively modeled using a cohort-aware machine learning framework. Public transcriptomic cohorts differ substantially in sample size, event rates and patient composition, all of which can bias model training toward larger or event-rich datasets^54^. To address this, we incorporated cohort-balanced weighting and optimized model selection using leave-one-cohort-out cross-validation, explicitly selecting models that generalized across independent cohorts. Under this framework, DeepSurv^43^ consistently outperformed conventional Cox regression in the validation cohorts. The improved performance likely reflects the combination of robust feature selection and cohort-aware model training, enabling MetaSig to maintain predictive performance across diverse patient populations and transcriptomic platforms.

MetaSig provided additional prognostic resolution within established TNBC subtype classifications. Although high-risk tumors were enriched within luminal androgen receptor and mesenchymal subtypes, prognostic differences were still observed within subtype groups, suggesting that subtype identity and clinical outcome capture related but non-equivalent dimensions of TNBC biology. Single-cell analyses further indicated that MetaSig reflects coordinated transcriptional programs across epithelial, endothelial, stromal and immune compartments rather than tumor-intrinsic programs alone. High-risk genes localized predominantly to tumor and stromal populations, whereas protective genes were enriched in adaptive immune cells. Together with the enrichment of hypoxia, glycolysis and extracellular matrix remodeling in high-risk tumors and interferon signaling in low-risk tumors, these findings support an ecosystem model of TNBC progression^55^. Consistent with this interpretation, regulatory network and CRISPR dependency analyses identified functionally relevant regulators underlying these transcriptional states. Integrating prognostic modeling with perturbational transcriptomics also illustrates how molecular classifiers can move beyond patient stratification to therapeutic hypothesis generation^56^. Drug prioritization against the Tahoe-100M atlas recovered established breast cancer therapeutics among the highest-ranking compounds, providing an internal validation of the framework, while also identifying additional agents predicted to reverse the transcriptional state of high-risk epithelial cells^46^. Although these predictions require functional validation, they demonstrate that reproducible prognostic programs may simultaneously define therapeutic vulnerabilities. This is particularly relevant for TNBC, where clinically actionable molecular targets remain limited despite extensive genomic characterization^2,57,58^.

Several limitations should be acknowledged. The analyses were performed retrospectively using publicly available cohorts with heterogeneous treatment regimens and follow-up, and prospective evaluation will ultimately be required before clinical implementation. Likewise, the therapeutic analyses are computational and should be considered hypothesis-generating until validated experimentally. Nevertheless, the consistent performance of MetaSig across independent cohorts, together with its concordance across bulk transcriptomic, single-cell, regulatory network and functional dependency analyses, suggests that the classifier captures biologically meaningful processes that extend beyond individual datasets.

In summary, this study demonstrates that the diversity of published TNBC prognostic gene signatures masks substantial convergence onto a relatively small number of conserved biological programs. By combining systematic cross-cohort benchmarking with independent transcriptome-wide biomarker discovery, machine-learning-based survival modeling and multi-layer biological characterization, we provide a framework that emphasizes reproducibility before prediction. More broadly, these findings suggest that clinically useful prognostic biomarkers will emerge not from increasingly complex prognostic gene signatures, but from molecular representations of biological processes that remain consistent across independent patient populations. To facilitate broad adoption and research utility, we provide a freely accessible web platform that enables users to calculate prognostic risk scores and risk-group classifications from normalized gene expression data https://metasig.igib.res.in.

## Methods

### Compilation of Prognostic Gene Signatures

To identify published prognostic signatures for TNBC, we systematically searched PubMed (https://pubmed.ncbi.nlm.nih.gov/) and Google Patents (https://patents.google.com/) using the terms “Triple Negative Breast Cancer Prognostic Gene Signature” OR “Biomarker”. Gene identifiers were standardized to approved HGNC symbols, with aliases and outdated identifiers resolved where possible. Eligible signatures were required to: (i) be developed for prognostic prediction in TNBC; (ii) be derived from transcriptomic data; (iii) contain at least three genes; and (iv) provide a complete and unambiguously mappable gene list. Signatures with unresolvable identifiers were excluded.

### Patient Cohorts and Data Processing

Transcriptomic and clinical data were obtained from six publicly available primary TNBC cohorts. Selection criteria were: (i) pre-treatment tumor samples obtained at diagnosis; and (ii) available survival data, including disease-free survival (DFS), recurrence-free survival (RFS), metastasis-free survival (MFS), or overall survival (OS), harmonized throughout as event-free survival (EFS) and OS. Discovery cohorts included three RNA-seq cohorts - TCGA-BRCA^6^, FUSCC^5^, and SCAN-B^10^ - and three microarray cohorts: METABRIC^7^, GSE58812^8^, and GSE21653^9^. Two independent cohorts - the Institut Jules Bordet cohort (JBordet)^59^ and GSE19615^60^ - served exclusively as external validation cohorts and were not used at any stage of gene selection, threshold optimization, or model training.

For TCGA, TPM values were retrieved using TCGAbiolinks (v2.36.0)^61^. FUSCC paired-end RNA-seq data were downloaded via SRA Toolkit (v2.10.9), quality-assessed with FastQC (v0.12.1) and MultiQC (v0.4), trimmed with Trimmomatic (v0.39; Phred <15), aligned to GRCh38 (Gencode v44) using STAR (v2.7.10b), and quantified as TPM using RSEM (v1.3.3)^5,62–65^. SCAN-B FPKM values were converted to TPM by within-sample normalization. METABRIC preprocessed log2-normalized values, GSE58812 MAS5-normalized values, and GSE21653 log2-ratio values were used as provided. For RNA-seq cohorts, TPM values were log2-transformed. Batch correction was applied to cohorts with available batch annotations (TCGA, FUSCC, METABRIC, SCAN-B) using ComBat from the sva R package (v3.56.0), applied independently of all clinical outcome variables^66^. Only genes shared across all six discovery cohorts were retained for downstream analysis.

TNBC status was defined by ER, PR, and HER2 negativity per ASCO/CAP guidelines^67,68^. PAM50 intrinsic sub-typing was performed using the genefu R package (v2.39.1)^15^. TNBCtype classification used the Vanderbilt TNBCtype framework^17^, refined to four stable subtypes^18^. Burstein TNBC sub-typing was performed by centroid-based Pearson correlation^19^.

### Signature Scoring and Quality Control

All 62 signatures were evaluated using the unweighted single-sample scoring methods ssGSEA and SingScore. For signatures with published coefficient-based formulas (n = 38), the original weighted risk scores were additionally calculated according to the corresponding publications. To enable uniform evaluation, all signatures were scored using two single-sample methods: ssGSEA (GSVA R package v2.2.1)^12^ and SingScore (singscore R package v1.28.1)^13^. For both ssGSEA and SingScore, genes reported as positively associated with poor prognosis were assigned to the up-regulated (high-risk) gene set, whereas genes associated with favorable prognosis were assigned to the down-regulated (low-risk) gene set. The final signature score was calculated as the up-regulated minus the down-regulated component, with signatures lacking a directional component assigned a value of zero for the absent gene set.

Signature quality was assessed using the signifinder R package (v1.10.0)^14^ via the evaluationSignPlot function, which evaluates gene coverage, zero-expression proportion, and correlation of scores with sequencing depth or zero-value percentage. These metrics were aggregated into a composite goodness score (0-100) per signature per cohort. In addition, to identify groups of biologically related prognostic signatures, consensus clustering was performed on the cohort-wise Z-score normalized ssGSEA signature scores using the ConsensusClusterPlus R package (v1.70.0)^69^, with consensus k-means clustering used to define robust signature clusters.

### Meta-Analysis Framework

An overview of the analytical workflow used for threshold harmonization, multivariable modeling, and meta-analysis is shown in **Supplementary Figure S1.** Multivariable and univariate survival analyses were conducted across all six discovery cohorts for both EFS and OS, using ssGSEA and SingScore stratifications, yielding four analysis combinations: ssGSEA-EFS, ssGSEA-OS, SingScore-EFS, and SingScore-OS. EFS and OS were analyzed independently because they reflect distinct clinical processes: EFS captures recurrence and disease progression, whereas OS additionally reflects post-relapse treatment, comorbidities, and competing causes of death^70,71^.

For each signature in each cohort, an optimal stratification threshold was determined using the maximally selected rank statistic (maxstat; R package v0.7.26)^72^, iterated 1,000 times over randomly sampled 50% patient subsets to ensure stability, with the median retained as the final threshold (Median Cohort Threshold). To account for inter-cohort variability, analyses were harmonized by applying each cohort-derived threshold to all six cohorts in turn, generating multiple sets of multivariable results per signature. Multivariable Cox models incorporated lymph node status, tumor size (T1 vs. T2-T4), and age (dichotomized at cohort-specific medians) as covariates, dynamically based on availability; GSE58812 models included age only. Meta-analysis across threshold sets used a random-effects model (meta R package v8.2.1)^73^. Signatures were retained only if they achieved consistent significance across all meta-analyses of different threshold sets. Complementary univariate meta-analyses and weighted risk score-based multivariable meta-analyses (for the 38 signatures with published weights) were also performed.

### Multi-Cohort Gene Discovery and Model Training

#### Gene-Level Survival Analysis and Derivation of Directionally Concordant Gene Sets

To derive a novel prognostic gene signature independent of previously published models, a gene-level survival analysis was performed across the six discovery TNBC transcriptomic cohorts, conducted separately for event-free survival (EFS) and overall survival (OS). Gene expression values were first Z-score normalized within each cohort, and all subsequent threshold estimation and feature selection steps were performed on these scaled values.

For each gene, an optimal expression threshold was determined using the maximally selected rank statistic (maxstat)^72^ approach- implemented via the maxstat R package (v0.7.26), iterated 1,000 times over randomly sampled 50% patient subsets to ensure stability, with the median threshold retained as the final threshold. Univariate Cox proportional hazards regression was then performed independently in each of the six cohorts. Because the discovery cohorts differed substantially in sample size and event rate, a gene’s apparent prognostic significance in a pooled analysis could be disproportionately driven by larger or event-rich cohorts rather than representing a truly reproducible biological association. To minimize this bias, prognostic evaluation was performed separately within each cohort, and cohort-balanced weighting based on sample size and event rate was applied throughout gene discovery and subsequent model training, ensuring that no single cohort dominated the overall assessment. A two-stage filtering strategy was applied: genes were required to reach statistical significance (P < 0.05) in all six cohorts, and then to exhibit a consistent direction of effect - uniformly elevated risk (HR > 1) or uniformly protective (HR < 1) - across all cohorts. Genes with discordant hazard ratio directions in any cohort were excluded regardless of significance. Genes satisfying both criteria were designated Directionally Concordant Gene Sets (DCGS), yielding two candidate sets: DCGS-EFS and DCGS-OS.

#### Within-Cohort Feature Refinement by Penalized and Stepwise Selection

To further distill the DCGS pools, two complementary feature selection procedures were applied independently within each of the six discovery cohorts: forward selection (FS) and least absolute shrinkage and selection operator (LASSO) selection (LS). LASSO-penalized Cox regression was performed using CoxnetSurvivalAnalysis (scikit-survival)^74^, with the optimal regularization strength selected by three-fold stratified cross-validation using concordance index as the optimization criterion; genes with non-zero coefficients at the optimal penalty were retained. Separately, FS was performed using Cox proportional hazards regression, iteratively adding the gene that most improved the mean cross-validated concordance index, with a minimum gain threshold of 0.005 required at each step to prevent inclusion of marginally informative features.

Genes identified by each procedure were aggregated across cohorts, and only those selected in at least four of the six discovery cohorts were retained-a consensus requirement consistent with the cross-cohort stringency applied during DCGS derivation. This yielded six gene configurations carried forward for model training: the original DCGS-EFS and DCGS-OS, together with their FS- and LS-derived subsets.

#### Expression Data Preparation for Model Training

For training the models - a parallel expression matrix was prepared specifically for the penalized Cox and DeepSurv model training procedures. Cross-cohort expression distributions were harmonized by applying Feature-Specific Quantile Normalization (FSQN) with the METABRIC cohort as the distributional reference, yielding a training matrix with consistent gene-level distributions across all six cohorts irrespective of the underlying platform^75^.

Validation cohort expression matrices were processed entirely independently of the discovery set. For each gene, scaling parameters - mean and standard deviation - were derived exclusively from the discovery matrix and applied to each validation cohort individually, ensuring that no information from validation patients influenced normalization parameters at any stage.

#### Cohort-Balanced Model Training

To prevent larger cohorts from disproportionately dominating model training, and to ensure that cohorts with high follow-up censoring contributed survival information proportional to their effective sample size, a two-component weighting strategy was applied at the cohort level.

A size-based weight was defined for each cohort *i* as:

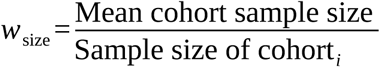

such that smaller cohorts received proportionally greater weight relative to the discovery mean. A complementary event-rate weight was defined as:

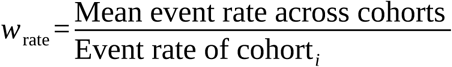

which up-weighted cohorts with lower observed event rates.

This component up-weighted cohorts with lower event rates - reflecting higher rates of right-censored observations - to offset the reduced per-patient survival information they contribute to likelihood-based training.

To avoid compounding inflation for cohorts that were simultaneously small and event-sparse, each component was independently capped at 3.0 prior to rescaling. Both components were then individually normalized to a mean of 1.0 across cohorts, and their mean was used to produce the final per-patient weight, denoted *w*:

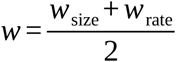

This additive formulation ensured that cohort size and event rate each contributed equally to the final weight without producing extreme values through their joint product. The resulting per-patient weight *w* was supplied directly to the CoxPH model via the weights_col argument in lifelines Python library^76^, and to the DeepSurv model as a sample weight tensor incorporated into the weighted Cox partial log-likelihood at each training step.

Two model architectures were trained on each gene configuration. A ridge-penalized Cox proportional hazards model (CoxPH) was fitted using CoxPHFitter from lifelines, with cohort weights and robust variance estimation. A DeepSurv model - a Cox proportional hazards deep neural network - was implemented in PyTorch (v≥2.0) following Katzman et al.^43,77^, comprising a fully connected multilayer perceptron that outputs a log-hazard score optimized via the weighted Cox partial log-likelihood. Training incorporated gradient clipping (L2 norm ≤ 1.0), best-epoch checkpointing, and early stopping after 20 epochs without improvement to ensure stable convergence.

Hyper-parameters for both architectures were optimized using a leave-one-cohort-out (LOCO) cross-validation framework, where each of the six discovery cohorts was withheld in turn as the validation set. This approach was preferred over random k-fold cross-validation as it directly evaluates cross-cohort generalization and prevents within-cohort data leakage. The CoxPH ridge penalizer was selected by grid search; DeepSurv architecture and regularization hyper-parameters were jointly optimized by Bayesian optimization using the Tree-structured Parzen Estimator in Optuna (v≥3.0)^78^. The configuration maximizing mean concordance index across LOCO folds was used to train a final model on the complete six-cohort matrix.

#### Performance Evaluation and Comparative Analysis

Prognostic discrimination was assessed using concordance index (C-index), AUROC at three years (AUROC₃), and AUROC at five years (AUROC₅), with 95% confidence intervals estimated by stratified bootstrapping over 300 replicates. For risk stratification, an optimal score threshold was determined per model by maximizing the log-rank statistic across discovery cohorts over a grid of 300 candidate threshold spanning the 10^th^-90^th^ percentile of the discovery score distribution, then applied without modification to all validation cohorts.

Discrimination metrics were computed across a unified set of scoring strategies applied consistently to both published and novel gene configurations. For all 62 compiled published signatures, four scoring approaches were evaluated in each cohort: ssGSEA-based enrichment scores, SingScore-based enrichment scores, and risk scores derived from CoxPH and DeepSurv models trained on each signature’s gene set. For the 38 published signatures for which explicit gene weights were available in the original publications, a weighted risk score was additionally computed per patient and evaluated by C-index, yielding a fifth scoring strategy for this subset. For the novel gene configurations-DCGS-EFS, DCGS-OS, and their forward selection (FS)- and LASSO selection (LS)-derived subsets-the same four strategies (ssGSEA, SingScore, CoxPH, and DeepSurv) were applied identically. This parallel evaluation across all scoring strategies and gene configurations enabled direct comparison of supervised machine learning approaches against single-sample enrichment methods, and benchmarking of novel signatures against the full landscape of previously published TNBC prognostic signatures, for both EFS and OS endpoints.

To identify reliably performing signatures across this comparative framework, a multi-criterion filter was applied: a model was considered reliable only if it achieved a mean discovery C-index ≥ 0.55, a train-to-validation performance gap ≤ 0.20, a validation-over-training excess ≤ 0.15, and C-index > 0.50 in at least four of the six discovery cohorts. Signatures failing any criterion were excluded from the final ranking.

#### Master Regulator Analysis

Master regulator (MR) analysis was performed to identify transcription factors (TFs) exhibiting differential activity between high-risk and low-risk patient groups using the ARACNe (Algorithm for the Reconstruction of Accurate Cellular Networks with Adaptive Partitioning)^79^ and VIPER^80^ (Virtual Inference of Protein-activity by Enriched Regulon analysis) frameworks. Gene regulatory network was constructed on the discovery cohort using the ARACNe-AP package^81^ with default parameters, following the protocol described by the authors (https://github.com/califano-lab/ARACNe-AP). A curated list of human TFs obtained from the DoRothEA R package (v1.18.0)^82^ was used as input for network inference. The generated ARACNe mutual information network was subsequently converted into a VIPER-compatible regulon object using the *aracne2regulon* function implemented in the viper R package (v1.40.0). Transcription factor protein activity scores were estimated for each sample using the viper function from the viper R package. Differential TF activity between high-risk and low-risk groups was assessed using t-test, with statistical significance evaluated against a null distribution generated through sample-label permutation. This was conducted using the *msviper* framework within the viper R package. In addition, leading-edge analysis, shadow analysis, and synergy analysis were performed according to the viper package documentation to further characterize regulatory dependencies and co-regulatory interactions.

#### Genetic Dependency and Drug-sensitivity Analysis

The genetic dependency of MetaSig-EFS, MetaSig-OS and their corresponding MRs was investigated using genome-wide CRISPR dependency data obtained from the DepMap portal^83^ (https://depmap.org/portal/download/). The Gene effect scores were retrieved for more than 1,300 cancer cell lines, including 19 TNBC cell lines from Cancer Cell Line Encyclopedia (CCLE)^84^. Gene effect scores below −0.5 were considered indicative of gene essentiality for cell survival, with more negative scores reflecting stronger cellular dependency on the corresponding gene. Conversely, a score of 0 indicated no detectable dependency or essentiality.

#### Single Cell Analysis

The breast cancer single-cell RNA-seq dataset GSE176078^44^ was downloaded from the Gene Expression Omnibus (GEO) database. Based on the associated study metadata, we selected 7 primary, treatment-naïve TNBC patients. Cell identities were assigned using the original cell annotations provided within the dataset, and normal epithelial cells were excluded prior to downstream analyses. Quality control filtering was applied to retain cells with more than 200 detected genes, more than 250 unique molecular identifier (UMI) count, and mitochondrial percentage below 10%. Data normalization, dimensionality reduction, and integration were performed using the Seurat R package (v5.4.0) to minimize inter-patient batch effects and technical variability^42,85–89^. Cell type-specific marker gene expression was assessed across the annotated populations, and only the genes shared with the discovery bulk RNA-seq dataset were retained. The final single-cell dataset used for the analysis comprised 25,972 cells and 16,045 genes spanning 16 distinct cell populations across epithelial, immune, stromal, and vascular compartments. Regulon activity at single-cell resolution was assessed using AUCell, which calculates an Area Under the Curve (AUC) score for each regulon in each cell based on the enrichment of regulon genes within the ranked gene expression profile^90^.

#### Identification of Risk-associated Cell Types

The Scissor R package (v2.0.0)^45^ was employed to integrate the discovery bulk RNA-seq dataset with single-cell RNA-seq data to identify cellular populations associated with the bulk-derived high-risk and low-risk transcriptional phenotypes. Briefly, Scissor first quantified the similarity between single-cell and bulk transcriptomic profiles by computing a correlation matrix. Subsequently, the cells associated with the risk phenotype were identified using the built-in logistic regression framework implemented in Scissor. Based on their association with the risk phenotype, cells were classified as Scissor+ (associated with the high-risk group), Scissor− (associated with the low-risk group), or background cells lacking a significant association.

The robustness and statistical significance of prognostic cell populations were assessed using 5-fold cross-validation implemented through Scissor’s *reliability.test()* function. This procedure evaluates whether the identified associations between bulk-derived phenotypes and single-cell transcriptional profiles are significantly stronger than expected by chance. The observed prediction performance was then compared against the randomized background using the area under the ROC curve (AUC). Associations between cell-type composition and risk groups were evaluated using Pearson’s Chi-squared Test implemented through the *chisq.test* function in the stats R package (v4.4.2) and the effect size was quantified through Cramér’s V calculated with the cramerV function from the rcompanion R package (v2.5.1). Differentially expressed genes (DEGs) between high-risk and low-risk associated cells were identified within each cell type using the FindMarkers function from the Seurat R package^85–89^. Consequently, functional enrichment analysis was subsequently performed on cell type-specific DEGs identified between high- and low-risk cell populations using Hallmark and C2 Reactome gene sets retrieved from the Molecular Signatures Database (MSigDB)^11^. Significantly enriched pathways (adjusted p-value<0.05) relevant to each gene set were selected for downstream visualisation.

#### Drug Repurposing using Tahoe-100M

Given the predominance of epithelial cells within the high-risk compartment, subsequent drug repurposing analyses were specifically focused on this population to identify therapeutic agents capable of reversing the high-risk transcriptional program. To this end, we developed a computational drug repurposing framework integrating single-cell differential expression disease signature with the Tahoe-100M perturbation dataset^46^ obtained from Hugging Face. Tahoe-100M comprises 100 million transcriptomic profiles characterizing the effects of 1,100 small-molecule compounds across 50 distinct cancer cell lines. The analysis was performed independently for MetaSig-EFS and MetaSig-OS disease signatures.

#### Disease Signature

Differentially expressed genes between high-risk and low-risk epithelial cells were used to define the disease query signature. The log2 fold change for each gene was used as the disease score, representing the direction and magnitude of transcriptional dysregulation. Genes with nominal p-value<0.05 were retained in the disease signature for downstream reversal scoring, yielding a gene set used both for instance-level Signature-Conditional Reversal Ordering Score (SCROS) and for computation of the final magnitude-weighted transcriptional reversal ratio.

#### Drug Perturbation Data

Drug-induced transcriptional profiles were obtained from the Tahoe-100M pseudo-bulk differential expression data. The analysis was restricted to HS-578T TNBC cell line to ensure disease-context specificity. Each experimental instance represented a unique combination of drug, concentration, and plate identifier. Instances were restricted to FDA-approved drugs as annotated in the Tahoe drug metadata. To reduce technical noise, drug names were retained in their original form as registered in the Tahoe dataset, preserving distinct formulation as separate entries.

#### Signal column selection

The Wald statistic was used as the primary drug signal, in preference to raw log2 fold change. The Wald statistic functions as a signal-to-noise ratio, automatically penalizing genes with large but unreliable fold changes due to high standard error, and operates on a Z-score-like scale that is more comparable across genes and instances.

Per-instance confidence filtering. Within each experimental instance, only genes meeting a statistical reliability threshold (Adjusted p-value < 0.1 where available, otherwise nominal p-value < 0.05) were retained. Instances yielding fewer than 10 confidently measured genes were excluded entirely. This prevents noisy low-quality experiments from contributing to downstream scoring.

#### Plate Normalisation

To correct for systematic plate-level batch effects, the median Wald statistic across all genes was computed per plate and subtracted from each observation on that plate, centering all plates at zero prior to any scoring.

#### Inverse-Variance Weighted Drug Profiles

Each drug was tested across multiple independent experimental instances in the HS-578T cell line (total 908 instances across all drugs). These replicates differ in concentration and plate, so a simple mean across them would treat a highly reproducible small effect the same as a large but erratic one. To address this, we computed a consensus expression profile for each drug using inverse-variance weighting (IVW), wherein, each gene measurement in each instance comes with an associated standard error (SE). A small SE means the measurement is precise and trustworthy; a large SE means it is noisy. IVW assigns each instance a weight equal to 1/SE², so precise measurements contribute more to the final consensus than imprecise ones. IVW profiles were computed exclusively over genes that had already passed the per-instance confidence filter (adjusted p < 0.1), ensuring that the gene universe used for scoring and for biological interpretation are identical. Drugs for which fewer than three HS-578T instances were available were excluded from downstream significance testing.

#### Signature-Conditional Reversal Ordering Score (SCROS)

The core per-instance scoring metric, SCROS, was developed to quantify how specifically a drug opposes the disease gene expression signature. To capture this, the Ordering Score (OS), a rank-based concordance statistic, is computed exclusively on the subset of genes where the drug is already confirmed to be acting against the disease direction, rather than across all genes globally. This makes the score conditional on reversal: a drug that broadly perturbs transcription without specifically opposing the disease signature cannot score highly.

For each drug instance and disease signature, the following procedure was applied:

1. Common gene set. Genes present in both the disease signature and the drug instance profile were identified. Instances with fewer than 10 common genes were assigned a score of zero.
2. Reversal masking. For each common gene, a reversal was declared if the product of its disease score and its drug Wald statistic was negative - meaning the two values carry opposite signs. Concretely, a gene up-regulated in disease (positive disease score) must be down-regulated by the drug (negative Wald stat), or vice versa, to count as reversed. Let:

- N_common = total number of genes present in both the disease signature and the drug instance profile.
- n_reversed = number of those genes where the drug Wald statistic has the opposite sign to the disease score, *i.e.,* the drug pushes that gene back toward its healthy expression level. Instances where n_reversed < 5 were assigned a score of zero.
3. Within-reversal OS. Among the reversed genes only, genes were ranked by the absolute magnitude of their disease score and drug Wald statistic respectively. For each reversed gene, two beta distribution-based p-values were computed: p_min, testing whether the gene ranked highly in both lists, and p_max, testing whether it ranked low in both. The smaller of the two was selected as the gene-level p-value and converted to a Z-score via the inverse cumulative distribution function of the standard normal distribution: Z = −Φ⁻¹(p). The ordering score (OS) was then computed as the sum of Z-scores exceeding the significance threshold(Z ≥ −Φ⁻¹(0.05) = 1.645), normalized by √n_reversed.
4. SCROS formulation. The final SCROS for instance *j* was:

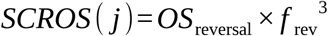

where 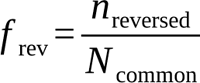is the fraction of common genes that the drug reverses. The cubic exponent was chosen to strongly penalize drugs with near-random reversal fractions relative to drugs with genuine signature specificity.

#### Robust Z-score Normalisation

SCROS values across all instances were normalized using a robust Z-score to reduce sensitivity to outliers. The global median and median absolute deviation (MAD) were computed and the Z-score was applied to all instances as:

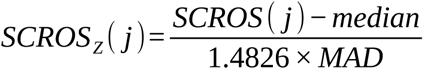

The MAD was scaled by the factor 1.4826, which makes it a consistent estimator of the standard deviation under normality, ensuring that the resulting Z-scores are interpretable on a standard scale. A floor of 10⁻⁶ applied to prevent division by zero.

#### Drug-Level Statistical Testing

For each drug, the distribution of SCROS Z-scores across its HS-578T instances (ranging from 3 to 6 per drug) was compared against the distribution across all other drugs’ instances using a one-sided Mann-Whitney U (Wilcoxon rank-sum) test with the alternative hypothesis that the drug’s scores are greater. Two safeguards were applied prior to testing: (i) a minimum of three instances was required to run the test, with drugs below this threshold assigned p = 1; (ii) a direction guard required the drug’s mean SCROS Z-score to exceed the background mean, otherwise p = 1 was assigned regardless of the U statistic. The MWU Z-score was computed from the U statistic and it was floored at zero, so no drug received a negative value.

#### Therapeutic Score

The final single-drug therapeutic score was defined as:

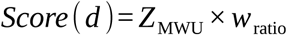

where *w*_ratio is the magnitude-weighted reversal ratio computed from the IVW drug profile against the significant disease DEGs (P < 0.05):

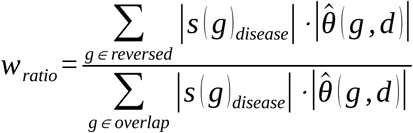

where s(g)disease is the disease score for gene g, *θ*^(^ *^g, d^* ^)^ is the IVW-combined Wald statistic for drug d at gene g, and the numerator sums only over genes classified as reversed (disease score and drug stat having opposite signs). While, the denominator sums over all genes present in both the IVW drug profile and the significant disease DEGs.

This formulation requires both statistical signal strength (MWU Z-score) and biological specificity (weighted reversal ratio) to be simultaneously high for a drug to achieve a high score. Genes with large disease dysregulation and large consistent drug response contribute proportionally more weight.

#### Permutation Validation and Artifact Filtering

To empirically assess pipeline specificity and identify drugs scoring high regardless of the disease signature content (potency artifacts), a row-shuffle permutation procedure was performed. In each of 10 permutations, the disease score and p-value columns of the epithelial DEG table were jointly shuffled across genes, destroying the gene-to-score mapping while preserving the marginal distributions of both scores and p-values. The full scoring pipeline was re-run on each permuted signature.

For each permutation, the top 10 ranked drugs were recorded. A permutation stability fraction was computed for each drug as the proportion of permutations in which it appeared in the top 10. Drugs with stability fraction ≥ 0.40 were classified as potency artifacts-their high scores were driven by general transcriptional activity in HS-578T rather than specific opposition of the disease signature and were excluded from the final drug list. The remaining drugs were re-ranked by their original therapeutic scores. This procedure provides empirical control over false positives at the pipeline level, complementing the formal statistical testing described above. Following artifact removal, normalized scores were recomputed within the filtered drug list.

#### Drug Combination Analysis

To identify complementary drug pairs, all pairwise combinations of the top 20 filtered drugs were evaluated. For each drug, a per-gene reversal profile was computed as *r* (*g, d*) = *−s_disease_* (*g*) *× θ*^^^ (*g, d*), where positive values indicate the drug opposes the disease direction at gene g. For each drug pair (A, B), a combined reversal profile was formed using a model: *r_combo_* (*g*) = *max* (*r _A_* (*g*) *, r _B_* (*g*)), meaning a gene is counted as reversed by the combination if either drug reverses it.

The combination weighted reversal ratio *(combo_w_ratio__)* was computed analogously to the single-drug *W_ratio_ratio__*. A synergy index was defined as the difference between the combination reversal ratio and the better of the two individual ratios. A complementarity index was defined as the fraction of reversed genes unique to one drug. Statistical significance of each combination was assessed using a one-sided MWU test comparing the pair’s per-gene reversal score vector against the pooled vectors of all other pairs, with FDR correction applied across all combinations tested. The combination therapeutic score was defined as:

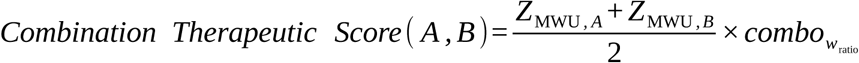

#### Positive Control Validation

Pipeline performance was assessed by verifying that established breast cancer therapeutics - including endocrine therapies (Anastrozole, Fulvestrant), CDK4/6 inhibitors (Palbociclib, Ribociclib, Abemaciclib), HER2-directed agents (Lapatinib, Neratinib), PARP inhibitors (Olaparib), cytotoxic chemotherapies (Paclitaxel, Docetaxel, Doxorubicin), and the mTOR inhibitor Everolimus - ranked within the top 50% of all scored drugs. Failure of known agents to achieve this threshold was treated as a signal for pipeline miscalibration.

## Statistical Analysis

All statistical analyses were performed using R (v4.5.1) and Python (v3.9.12). Pearson’s correlation coefficient was calculated to assess relationships between continuous variables. A p-value of < 0.05 was considered statistically significant unless otherwise stated. Multiple testing correction was performed using the false discovery rate (FDR) method where applicable.

Signature quality metrics, including gene coverage, zero value percentage, and correlation with sequencing depth, were evaluated using the signifinder R package to assess robustness of signatures for downstream analysis.

All survival analyses were based on time-to-event data, with time measured from the date of cancer diagnosis to the occurrence of an event of interest or last follow-up (censoring). Survival curves were estimated using the Kaplan-Meier method and compared using the log-rank test. Univariate and multivariable Cox proportional hazards regression models were fitted using the survival R package (v3.8.3)^91^ to assess associations between gene signatures and survival outcomes. Hazard ratios (HRs) with 95% confidence intervals (CIs) were reported. The proportional hazards assumption was assessed using Schoenfeld residuals.

Meta-analyses across cohorts were performed using the meta R package (v8.2.1) with random-effects models. Concordance index (C-index) was calculated to evaluate the discriminative ability of prognostic signatures. Statistical significance in Cox models was determined using the Wald test.

Functional enrichment analyses were conducted using the clusterProfiler R package (v4.14.6)^92^. Multiple testing correction was performed using the Benjamini-Hochberg false discovery rate (FDR) method, and pathways with an adjusted *p*-value (FDR) < 0.05 were considered statistically significant.

## Data Availability

All transcriptomic and clinical data used in this study are publicly available. RNA sequencing and clinical data for TCGA-BRCA were retrieved from the Genomic Data Commons portal (https://portal.gdc.cancer.gov), and METABRIC data were obtained from the cBioPortal for Cancer Genomics (https://www.cbioportal.org). SCAN-B cohort data were downloaded from Mendeley Data (https://data.mendeley.com/datasets/yzxtxn4nmd). FUSCC cohort RNA-sequencing data were obtained from the Sequence Read Archive (SRA) under accession SRP157974. Microarray data for GSE58812 and GSE21653 were retrieved from the NCBI Gene Expression Omnibus (GEO, https://www.ncbi.nlm.nih.gov/geo) under their respective accession numbers. The Institut Jules Bordet (JBordet) validation cohort data were obtained from Zenodo (https://doi.org/10.5281/zenodo.8135721), as reported in the original publication. The GSE19615 validation cohort was retrieved from GEO under its respective accession number. Single-cell RNA-seq data were obtained from GEO under accession GSE176078. Tahoe-100M perturbation differential expression data were retrieved from Hugging Face (https://huggingface.co/datasets/tahoebio/Tahoe-100M). All clinical and survival annotations were obtained from the same public sources as described in the original publications for each cohort. No new data were generated in this study.

The supplementary tables associated with this study are publicly available on Zenodo at: https://zenodo.org/records/21096162

The interactive web tool for signature score calculation is accessible at https://metasig.igib.res.in.

## Code Availability

The scripts and processed datasets used for data preprocessing, model development, statistical analyses, and figure generation will be made publicly available through a GitHub repository and archived in Zenodo upon publication of this manuscript. The repository link and DOI will be added to the final published version.

## Acknowledgments

LD and MS acknowledges CSIR-IGIB for support. SJ acknowledges the University Grants Commission (UGC) for fellowship support. DY acknowledges CSIR for fellowship support. SB acknowledges DBT/Wellcome Trust India Alliance Early Career Fellowship (ECF) (IA/I/22/1/506773) for funding support. LD, SJ, DY and SB gratefully acknowledge the Academy of Scientific and Innovative Research (AcSIR), Ghaziabad, India, for academic support. We acknowledge the support from CSIR-IGIB for infrastructure and its IT department for computing facilities

## Authors Contributions

SB conceived the study and led the overall project design. LD, MS, SJ and DY compiled published TNBC prognostic signatures and interpretation of results. SB, LD and MS carried out the multi-cohort data processing and survival analyses, development of the prognostic classifier and implementation of the prediction tool. SJ conducted the single-cell transcriptomic analyses. The manuscript was written by SB, LD, MS and SJ. All authors contributed to its editing and reviewed the final version.

