## Supplementary Figures for "Systematic Meta-Analysis of Published Transcriptomic Prognostic Signatures and Development of a Robust Multi-Cohort Prognostic Classifier for Triple-Negative Breast Cancer"

### Table of content of supplementary figures


| ***Supplementary Figure*** | ***Title*** |
|---|---|
| *Supplementary Figure S1* | *Overview of the study workflow* |
| *Supplementary Figure S2* | *Pairwise gene and pathway overlap among 62 published TNBC prognostic signatures* |
| *Supplementary Figure S3* | *Batch correction effectively reduces technical variance across TNBC discovery cohorts* |
| *Supplementary Figure S4* | *Harmonization of signature scores across discovery cohorts using ssGSEA and SingScore* |
| *Supplementary Figure S5* | *Univariate cross-threshold analysis reveals broader prognostic associations across cohorts and scoring methods* |
| *Supplementary Figure S6* | *Cross-threshold meta-analysis identifies prognostic signatures with consistent survival associations across independent TNBC cohorts* |
| *Supplementary Figure S7* | *Weighted risk score meta-analysis confirms prognostic associations in signatures with published scoring models* |
| *Supplementary Figure S8* | *Development and validation of MetaSig-OS, a multi-cohort prognostic classifier for TNBC* |
| *Supplementary Figure S9* | *MetaSig-EFS stratifies patients into distinct risk groups across established TNBC molecular subtype classifications* |
| *Supplementary Figure S10* | *MetaSig-OS stratifies patients into distinct risk groups across established TNBC molecular subtype classifications* |
| *Supplementary Figure S11* | *Pathway enrichment associated with MetaSig risk groups within TNBC molecular subtypes* |
| *Supplementary Figure S12* | *MetaSig-EFS and MetaSig-OS stratify patients independently of clinical variables* |
| *Supplementary Figure S13* | *Batch integration and cell type-specific marker expression in the TNBC single-cell dataset* |
| *Supplementary Figure S14* | *Upstream Regulatory Architecture of MetaSig-OS* |
| *Supplementary Figure S15* | *Risk-Associated Cell States and Candidate Therapeutics for MetaSig-OS* |
| *Supplementary Figure S16* | *Positive Control Validation for Drug Repurposing* |

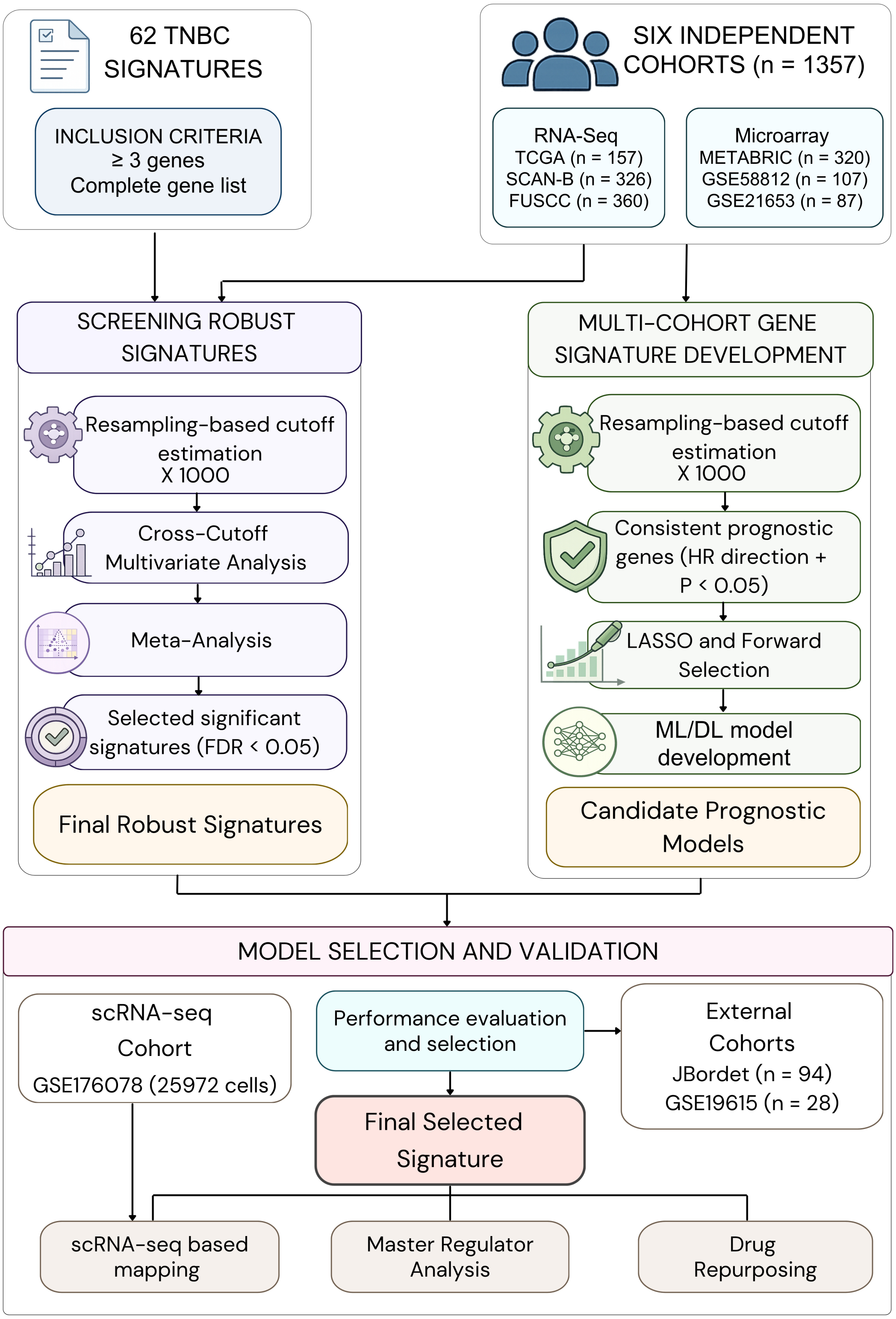


**Supplementary Figure S1. Overview of the study workflow.**
Published TNBC prognostic signatures were systematically collected and evaluated across six independent transcriptomic cohorts using a multi-cohort survival analysis framework. In parallel, a novel prognostic signature was developed through cross-cohort gene selection and machine learning-based modeling. The

final models were validated in external cohorts and further characterized using single-cell transcriptomic analysis, master regulator inference, and drug repurposing approaches.

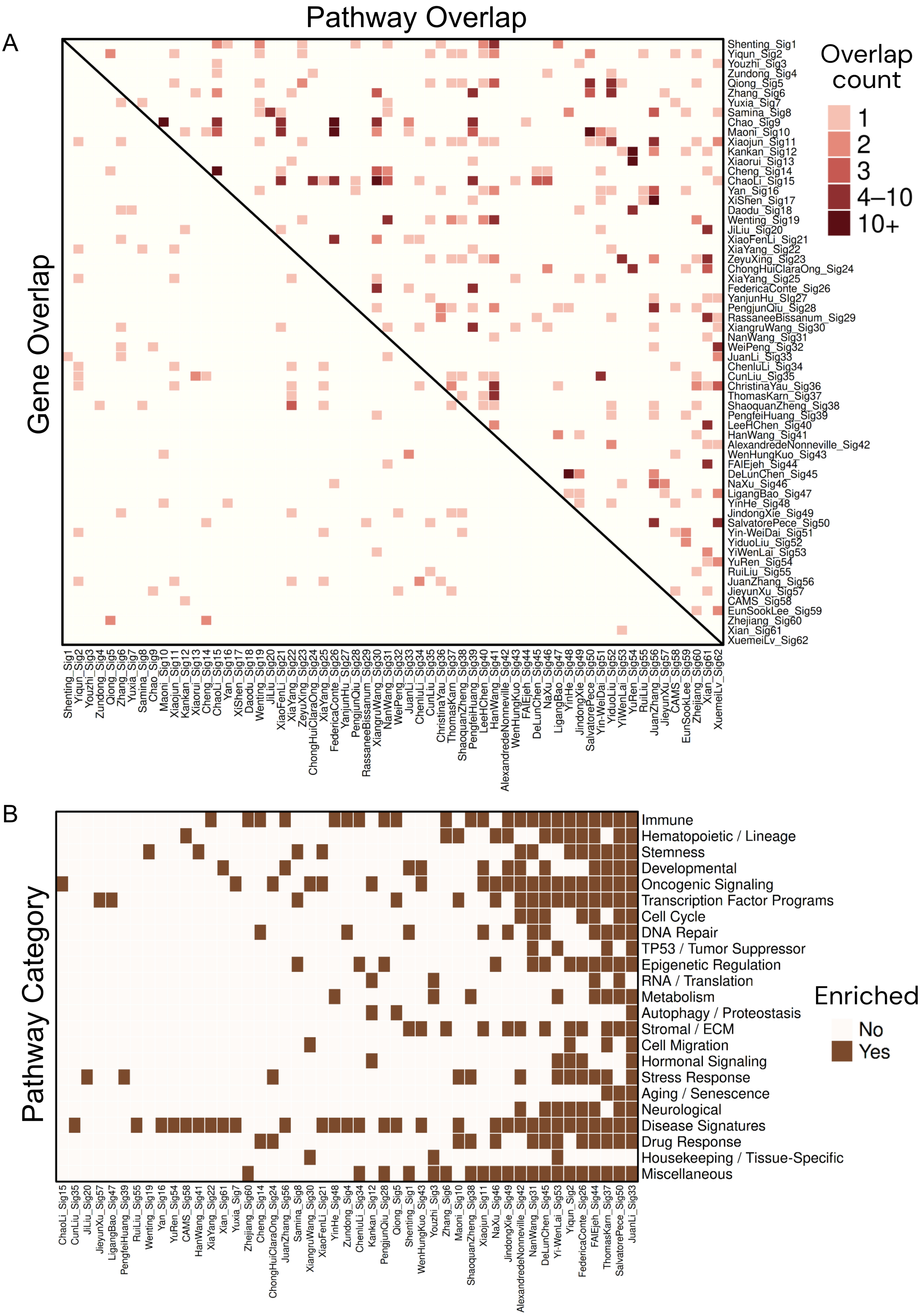


**Supplementary Figure S2. Pairwise gene and pathway overlap among 62 published TNBC prognostic signatures.**

A) Heatmap displaying pairwise overlap in gene composition (lower triangle) and enriched biological pathways (upper triangle) across 62 published TNBC prognostic signatures. Color intensity reflects the number of shared genes or pathways between each signature pair, ranging from 1 (light pink) to 10 or more (dark red), as indicated in the legend. Despite limited gene-level overlap across most signature pairs, substantially greater convergence is observed at the pathway level, with several signature clusters sharing common biological processes. Darker blocks along the pathway triangle indicate groups of signatures capturing overlapping transcriptional programs, suggesting that independently derived models frequently converge on shared biological mechanisms underlying TNBC prognosis. B) Enrichment of broad biological pathway categories across the 62 published TNBC prognostic signatures. Filled squares indicate significant enrichment of at least one pathway within the corresponding category. Although individual signatures share few genes, many converge on common biological programs, including immune, metabolic, stromal, and oncogenic processes.

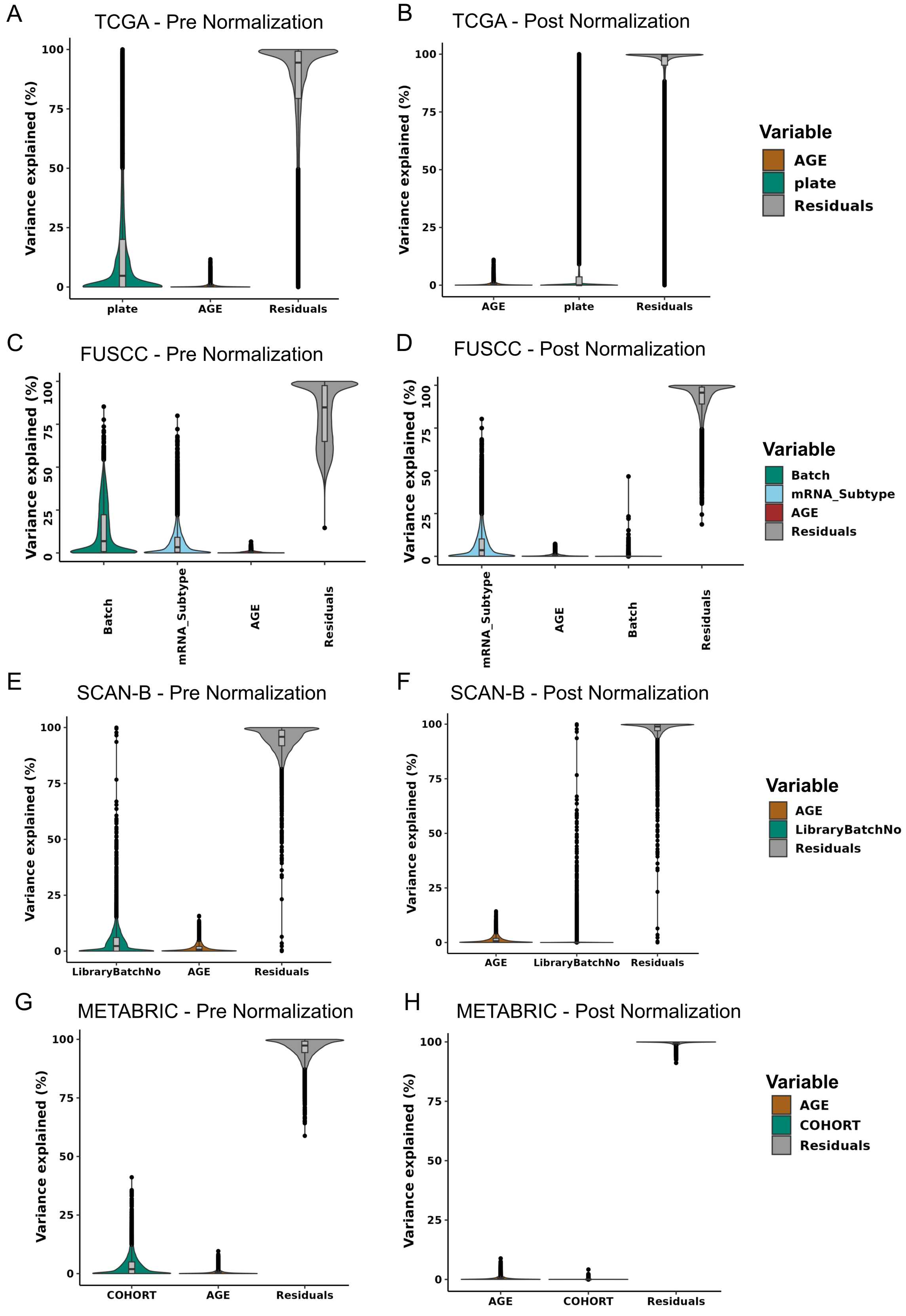


**Supplementary Figure S3. Batch correction effectively reduces technical variance across TNBC discovery cohorts.**

Variance Partition violin plots showing the percentage of gene expression variance explained by technical and biological variables before (left panels) and after (right panels) batch correction for TCGA (A, B), FUSCC

(C, D), SCAN-B (E, F), and METABRIC (G, H). Each violin represents the distribution of variance explained across all genes for the indicated variable. Batch-associated variables including sequencing plate (TCGA), batch (FUSCC), and library batch number (SCAN-B) showed substantially reduced variance contribution following ComBat-based normalization, while residual variance remained the dominant source of variation post-correction. Age and molecular subtype contributions remained largely stable across normalization steps, confirming that biological signal was preserved during batch correction.

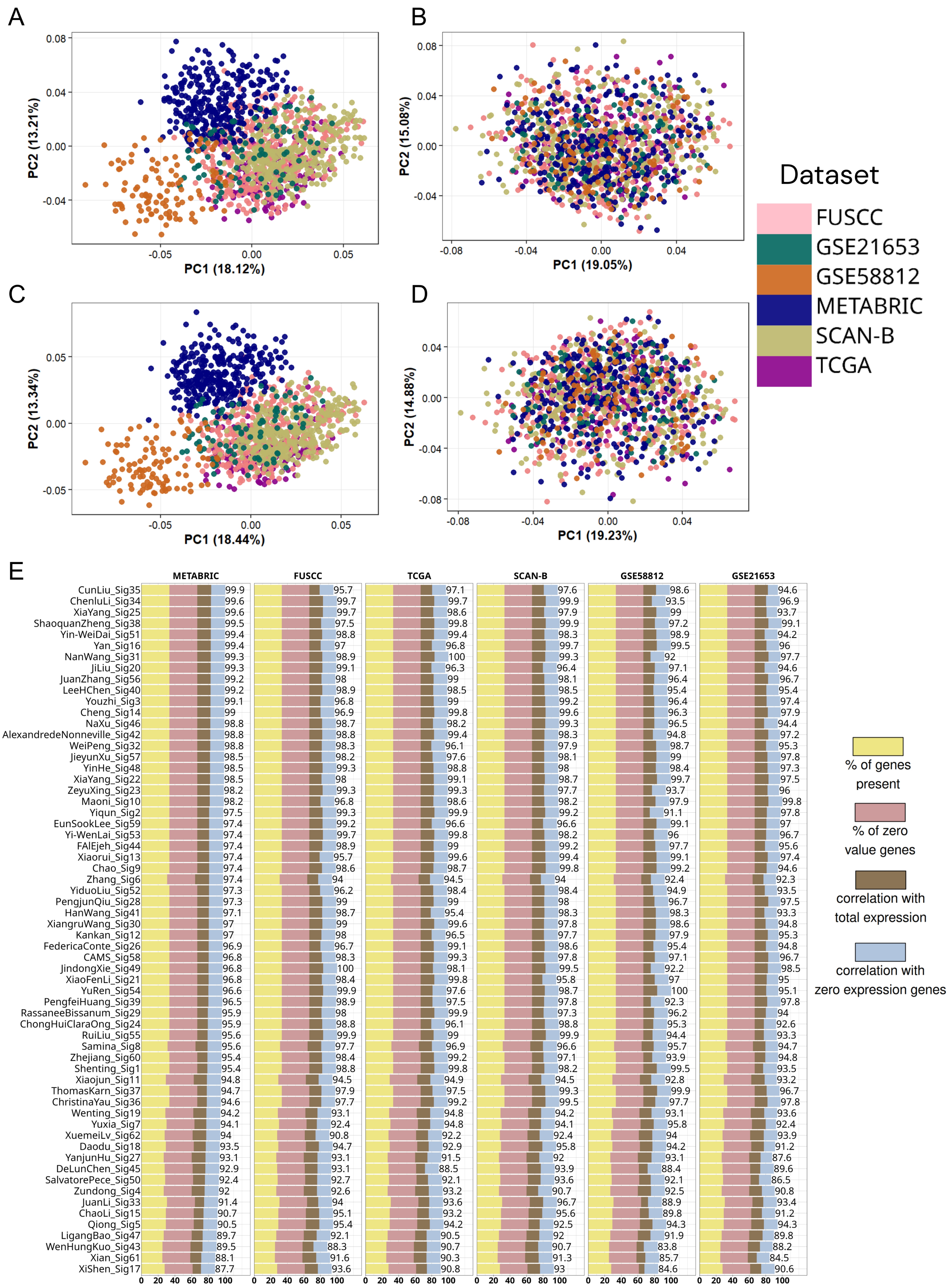


**Supplementary Figure S4. Harmonization of signature scores across discovery cohorts using ssGSEA and SingScore.**

PCA plots illustrating the distribution of per-patient signature scores across six TNBC discovery cohorts before (A, C) and after (B, D) within-cohort z-score normalization, computed using ssGSEA (A, B) and SingScore (C, D). Each point represents an individual patient colored by dataset. Prior to normalization, samples clustered by cohort, reflecting dataset-specific score distributions driven by differences in expression platforms and patient composition. Following z-score normalization within each cohort, samples were well-mixed across datasets in both scoring methods, confirming that score harmonization effectively removes cohort-specific distributional effects while preserving the relative ranking of patients within each dataset. Panel E displays signature quality metrics for all 62 published TNBC prognostic signatures across six cohorts as assessed using the signifinder package, including percentage of signature genes present, percentage of zero-value genes, correlation with total expression, and correlation with zero-expression genes. All signatures demonstrated high gene coverage and robustness across cohorts, supporting their suitability for standardized cross-cohort comparative analysis.

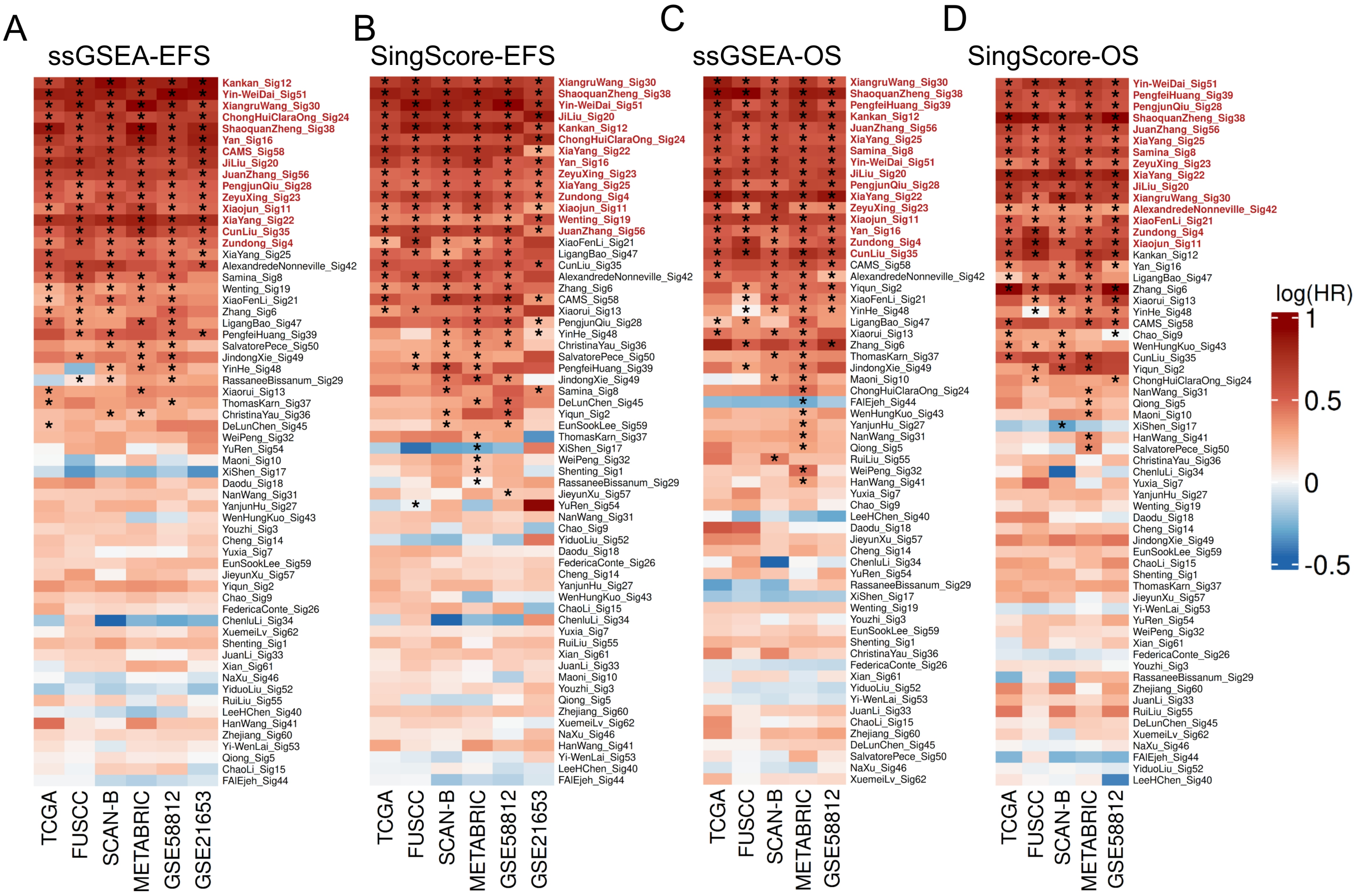


**Supplementary Figure S5. Univariate cross-threshold analysis reveals broader prognostic associations across cohorts and scoring methods.**

Heatmaps displaying hazard ratios (log scale) from univariate Cox regression across six discovery cohorts for 62 published TNBC prognostic signatures, evaluated under cross- threshold harmonization using ssGSEA-EFS (A), SingScore-EFS (B), ssGSEA-OS (C), and SingScore-OS (D). Each row represents a signature and each column a cohort, with color reflecting the direction and magnitude of the association (red = higher risk, blue = lower risk). Black dots indicate signatures reaching statistical significance within individual cohorts. Compared to the multivariable meta-analysis, univariate cross- threshold analysis identified a larger subset of signatures with consistent directional associations across cohorts, reflecting the contribution of clinical covariates in attenuating some prognostic signals in adjusted models.

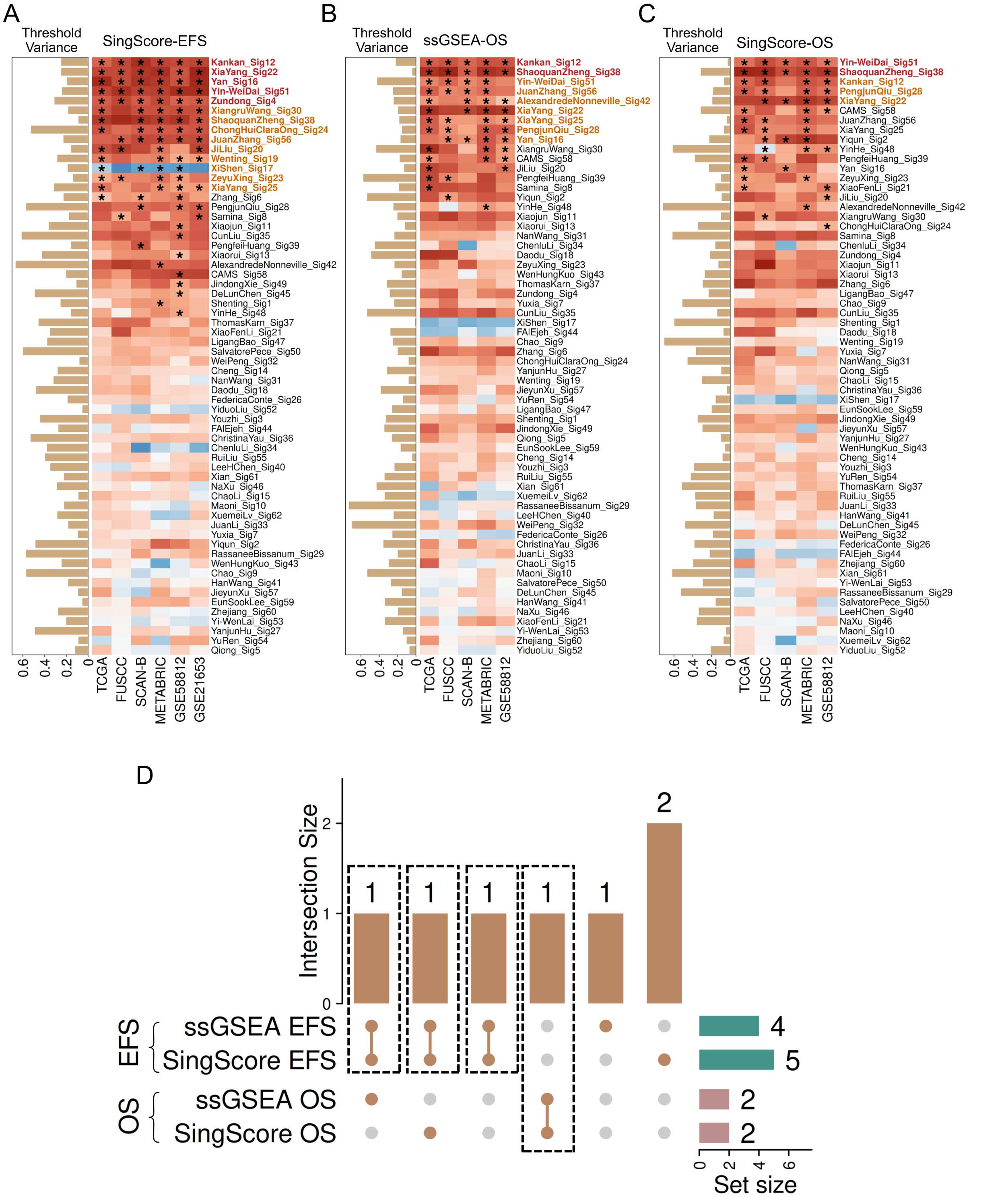


**Supplementary Figure S6. Cross-threshold meta-analysis identifies prognostic signatures with consistent survival associations across independent TNBC cohorts.**

Heatmaps displaying hazard ratios (log scale) from multivariable Cox regression meta-analysis for 62 published TNBC prognostic signatures across six discovery cohorts, evaluated using SingScore-EFS (A), ssGSEA-OS (B), and SingScore-OS (C). Each row represents a signature and each column a cohort, with color reflecting the direction and magnitude of the hazard ratio (red = higher risk, blue = lower risk). The variance bar on the left of each heatmap indicates cross-cohort variability in signature thresholds. Black dots denote signatures reaching statistical significance (FDR < 0.05). Signatures highlighted in colored text were consistently significant across all cross-threshold meta-analyses within the respective scoring and endpoint combination. (D) UpSet plot showing the intersection of consistently significant signatures across all four scoring-endpoint combinations: ssGSEA-EFS, SingScore-EFS, ssGSEA-OS, and SingScore-OS. Set sizes are shown on the left. Three signatures, Kankan_Sig12, Yan_Sig16 and Yin-WeiDai_Sig51, were consistently significant across both EFS scoring methods, while one signature : ShaoquanZheng_Sig38, was reproducibly significant for OS. The boxed intersections highlight signatures meeting significance criteria across multiple combinations, representing the most robustly validated prognostic models among all 62 evaluated signatures.

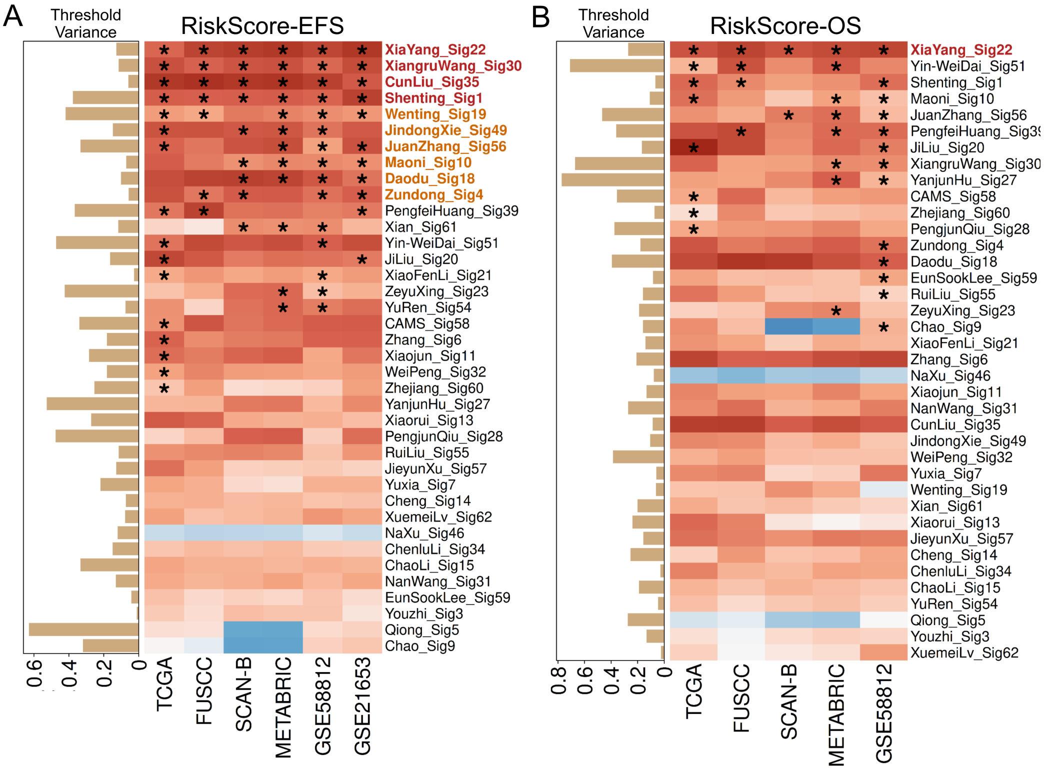


**Supplementary Figure S7. Weighted risk score meta-analysis confirms prognostic associations in signatures with published scoring models.**

For the 38 signatures with explicitly defined weighted risk scoring models, per-patient risk scores were computed using published gene-specific coefficients and patients were stratified at the dataset-specific median. Heatmaps show multivariable Cox hazard ratios (log scale) for EFS **(A)** and OS **(B)** across six cohorts, with asterisks denoting cohort-level significance and variance bars reflecting cross-cohort variability. Signatures in red were consistently significant for all datasets; those in orange were significant for at least 5 datasets.

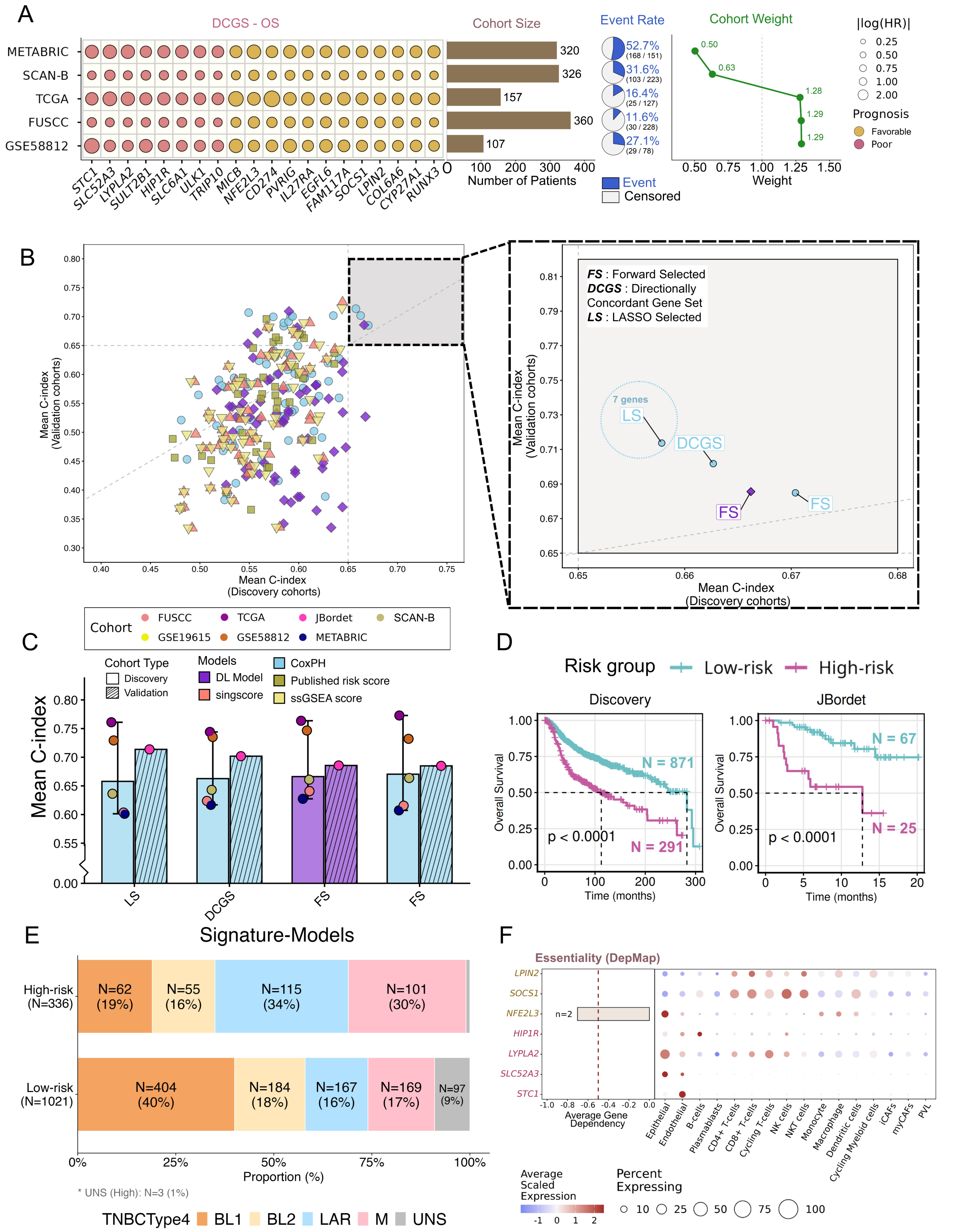

**Supplementary Figure S8 : Development and validation of MetaSig-OS, a multi-cohort prognostic classifier for TNBC**

A) Identification of the Directionally Concordant Gene Set (DCGS-OS) across the six discovery cohorts and the cohort-balanced weighting scheme used for model training. Dot size represents the pooled hazard ratio

(HR) magnitude and color indicates prognostic direction. The accompanying panel summarizes cohort characteristics, including sample size, event rate, and the relative cohort weights assigned during model development to balance differences in cohort size and outcome frequency. C) Performance of prognostic modeling strategies comparing published signatures, DCGS gene sets, and reduced models using Cox proportional hazards (CoxPH) and DeepSurv. Mean C-index in discovery cohorts is plotted against external validation cohorts; the inset highlights the best-performing models. MetaSig-OS (Lasso Selected DCGS-OS trained with CoxPH) achieved the highest overall performance. D) Mean C-index of the top-performing models across individual discovery and validation cohorts. Bars indicate discovery performance, striped bars indicate validation performance, and points represent cohort-specific C-indices. E) Kaplan-Meier curves showing overall survival stratified by MetaSig-OS risk groups in the combined discovery cohort and the independent validation cohort (JBordet). Risk groups were defined using the discovery cohort threshold (top 25% high-risk) and applied unchanged to the validation cohort. F) Distribution of TNBCtype-4 molecular subtypes across MetaSig-OS low- and high-risk groups. G) DepMap mean gene dependency scores and single-cell expression of MetaSig-OS genes across cell types in the TNBC tumor microenvironment.

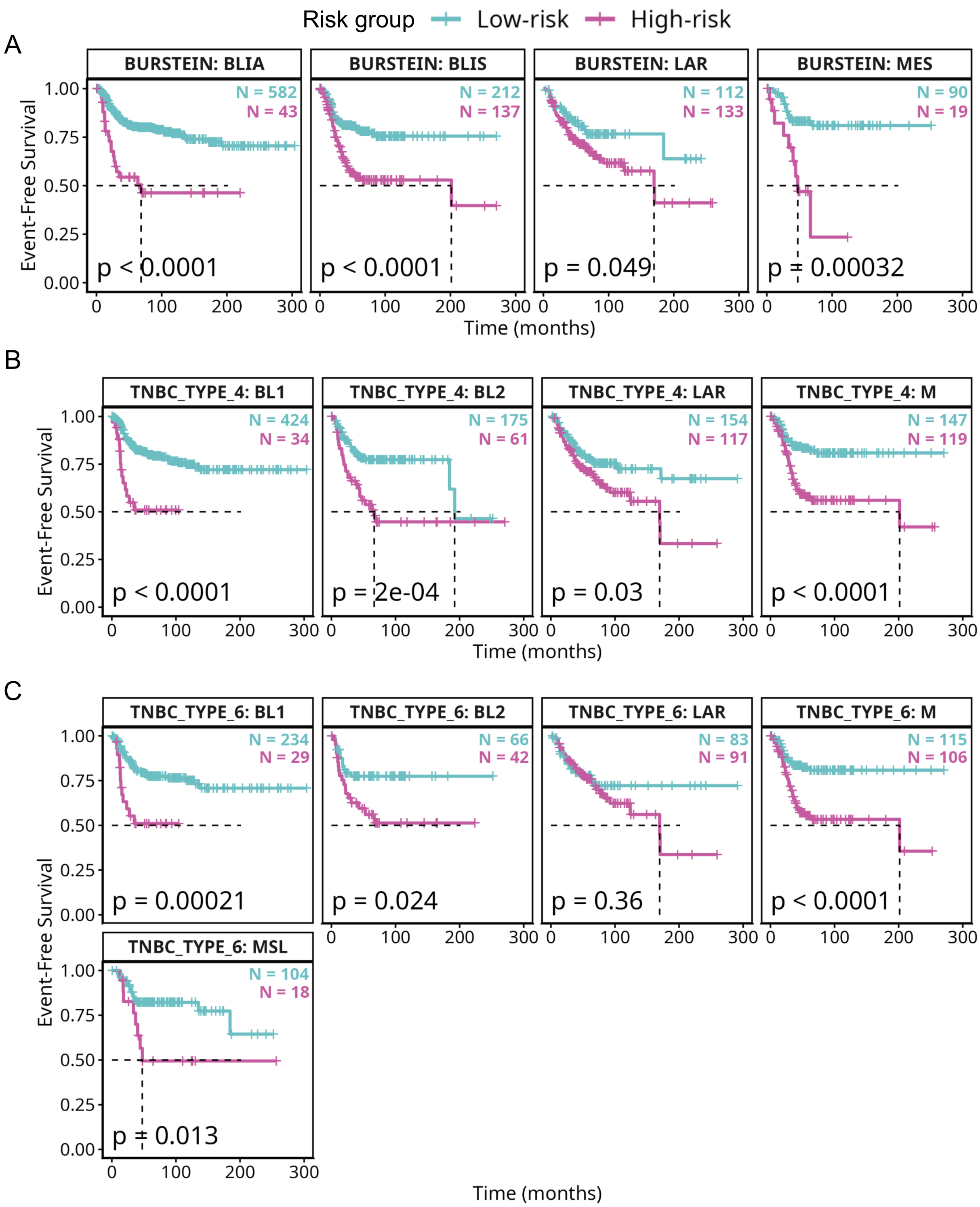


**Supplementary Figure S9. MetaSig-EFS stratifies patients into distinct risk groups across established TNBC molecular subtype classifications.**

Kaplan-Meier survival curves demonstrating prognostic separation by MetaSig-EFS within molecularly defined TNBC subgroups. High-risk (pink) and low-risk (cyan) groups were defined by MetaSig-EFS risk scores. A) Within Burstein subtypes, significant risk stratification was observed in BLIA ($p < 0.0001$), BLIS ($p < 0.0001$), LAR ($p = 0.049$), and MES ($p = 0.00032$) subgroups. B) Within TNBCtype-4 subtypes, MetaSig-EFS significantly separated high- and low-risk patients in BL1 ($p < 0.0001$), BL2 ($p = 2e-04$), LAR ($p = 0.03$), and M ($p < 0.0001$) subgroups. C) Within TNBCtype-6 subtypes, significant stratification was achieved in BL1 ($p = 0.00021$), BL2 ($p = 0.024$), M ($p < 0.0001$), and MSL ($p = 0.013$) subgroups, with a non-significant trend observed in LAR ($p = 0.36$).

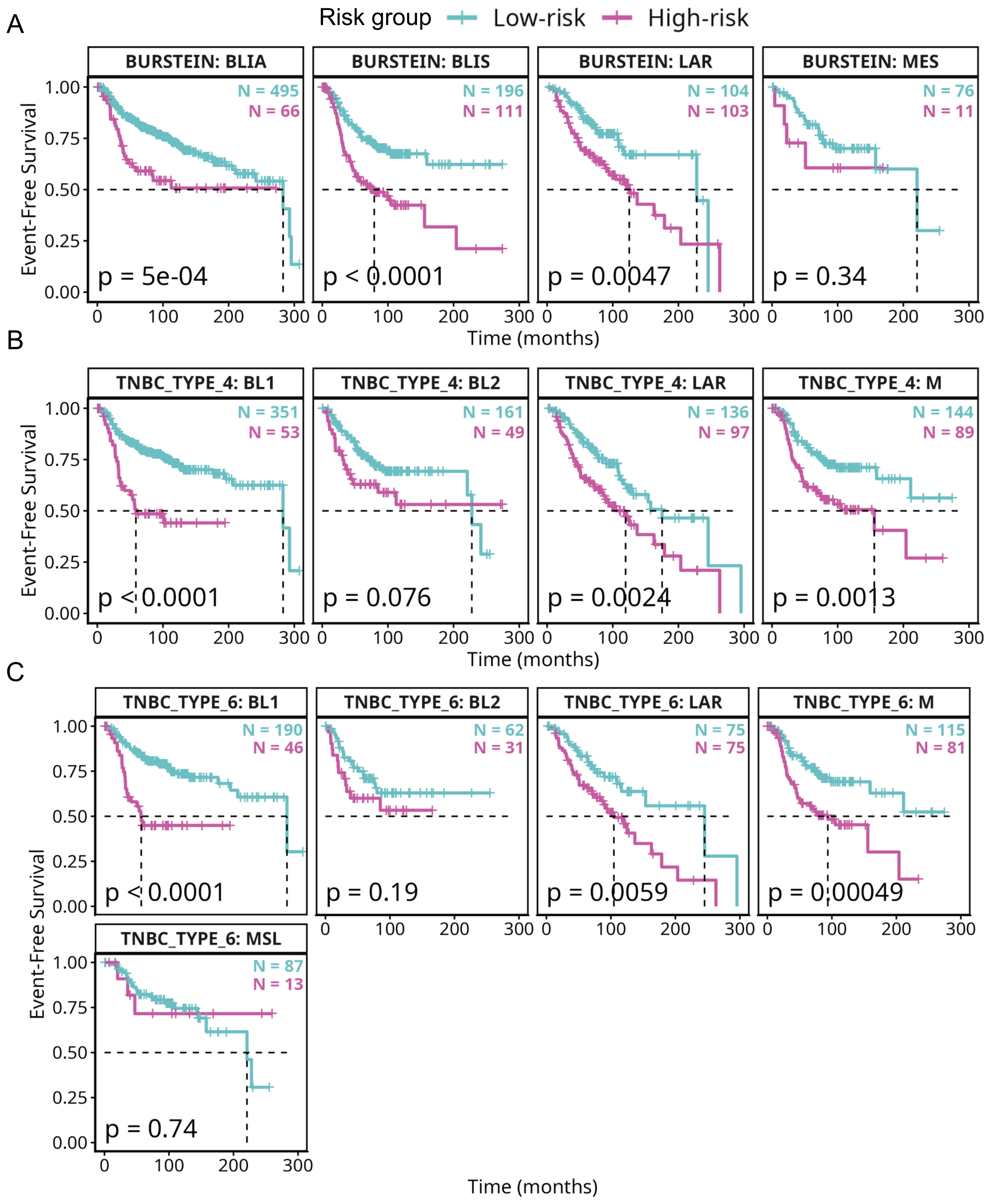


**Supplementary Figure S10.** MetaSig-OS stratifies patients into distinct risk groups across established TNBC molecular subtype classifications.

Kaplan-Meier survival curves demonstrating prognostic separation by MetaSig-OS within molecularly defined TNBC subgroups. High-risk (pink) and low-risk (cyan) groups were defined by MetaSig-OS risk scores. A) Within Burstein subtypes, significant risk stratification was observed across all four subgroups including BLIA ($p = 5e-04$ ), BLIS ($p < 0.0001$), LAR ($p = 0.0047$), and MES ($p = 0.34$). B) Within TNBCtype-4 subtypes, MetaSig-OS significantly separated high- and low-risk patients in BL1 ($p < 0.0001$), BL2 ($p = 0.076$), LAR ($p = 0.0024$), and M ($p = 0.0013$) subgroups. C) Within TNBCtype-6 subtypes, significant stratification was achieved in BL1 ($p < 0.0001$), BL2 ($p = 0.19$), LAR ($p = 0.0059$), and M ($p = 0.00049$) subgroups, with a non-significant trend in MSL ($p = 0.74$).

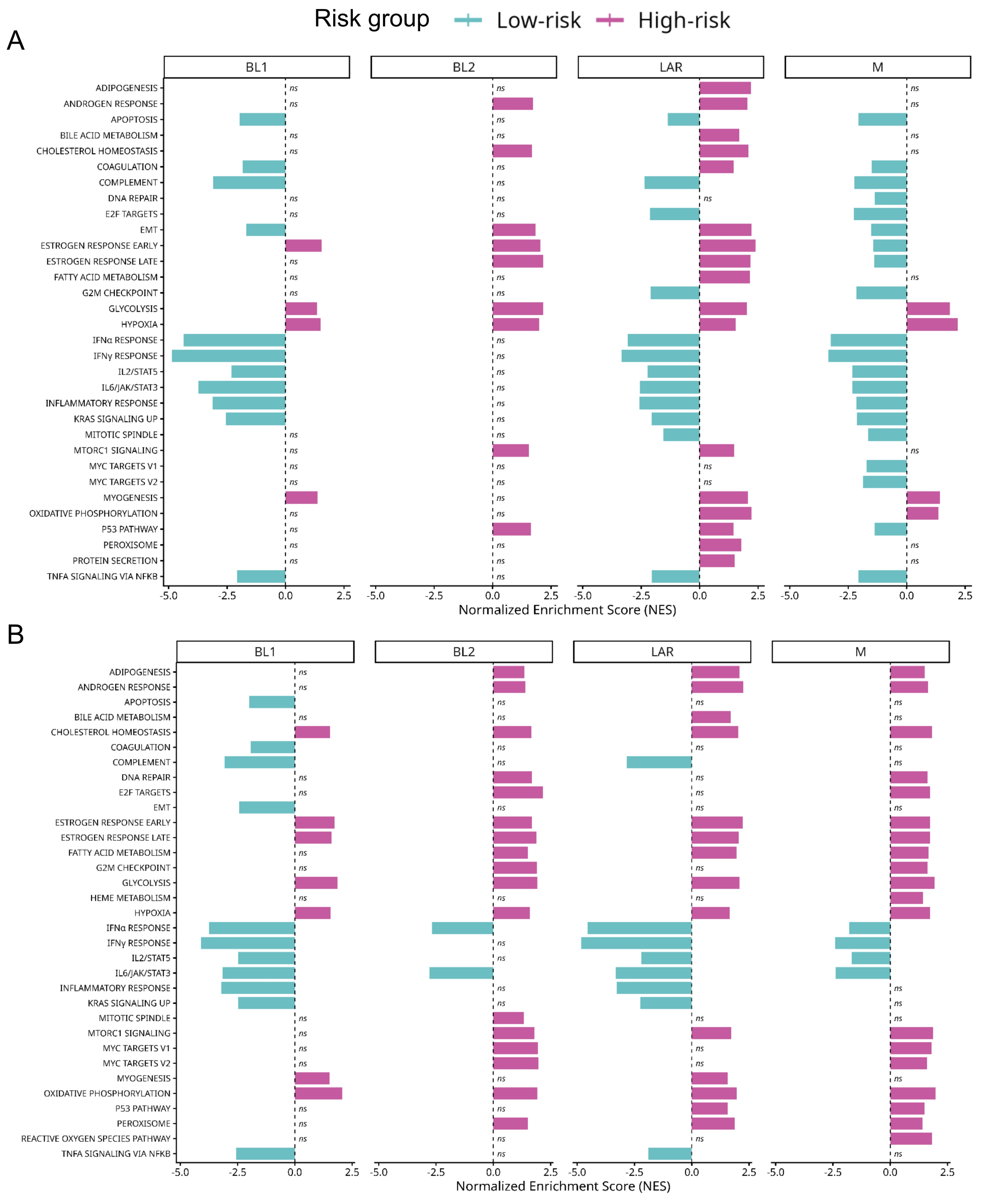


**Supplementary Figure S11. Pathway enrichment associated with MetaSig risk groups within TNBC molecular subtypes.**
A) MetaSig-EFS B) MetaSig-OS. Pathways are differentially enriched between low-risk and high-risk tumors within BL1, BL2, LAR, and M subtypes. Positive normalized enrichment scores (NES) indicate enrichment in

high-risk tumors, whereas negative NES values indicate enrichment in low-risk tumors. Statistical significance is indicated on the plots.

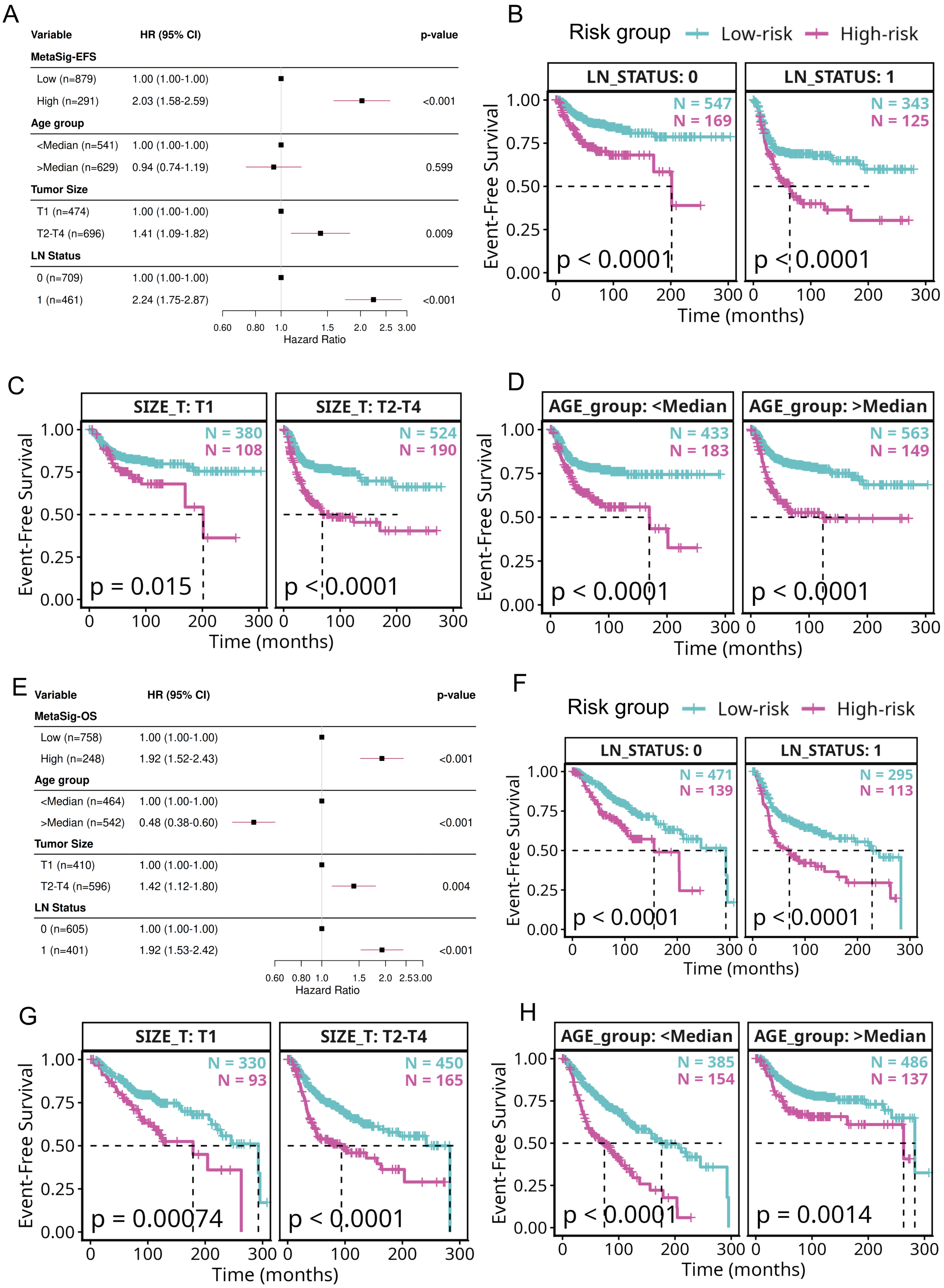


**Supplementary Figure S12. MetaSig-EFS and MetaSig-OS stratify patients independently of clinical variables.**

A) Forest plot from multivariable Cox regression analysis adjusting for age, tumor size, and lymph node status across the combined discovery cohorts. MetaSig-EFS high-risk group was independently associated

with significantly worse event-free survival (HR = 2.03, 95% CI: 1.58-2.59, $p < 0.001$), as were T2-T4 tumor size (HR = 1.41, p = 0.009) and node-positive status (HR = 2.24, $p < 0.001$). Kaplan-Meier survival curves demonstrating the prognostic performance of MetaSig-EFS (B-D) and MetaSig-OS (F-H) risk stratification within clinically defined patient subgroups. High-risk (pink) and low-risk (cyan) groups were defined by MetaSig-EFS and MetaSig-OS scores respectively. For MetaSig-EFS, significant separation between high- and low-risk patients was observed within node-negative (LN STATUS: 0, $p < 0.0001$) and node-positive (LN STATUS: 1, $p < 0.0001$) subgroups (B), within T1 (p = 0.015) and T2-T4 ($p < 0.0001$) tumor size categories (C), and within both below-median ($p < 0.0001$) and above-median ($p < 0.0001$) age groups (D).
E) Forest plot from multivariable Cox regression confirming independent prognostic value of MetaSig-OS after adjustment for age, tumor size, and lymph node status (HR = 1.92, 95% CI: 1.52 - 2.43, $p < 0.001$) for MetaSig-OS MetaSig-OS also maintained significant risk stratification within node-negative (p = 0.0011) and node-positive ($p < 0.0001$) subgroups (F), T1 (p = 0.029) and T2-T4 ($p < 0.0001$) tumor size categories (G), and below-median ($p < 0.0001$) and above-median (p = 5e-04) age groups (H). The consistent and significant separation across all clinical strata demonstrates that both signatures provide prognostic information independent of lymph node status, tumor size, and patient age, supporting their utility as independent biomarkers for risk stratification in TNBC.

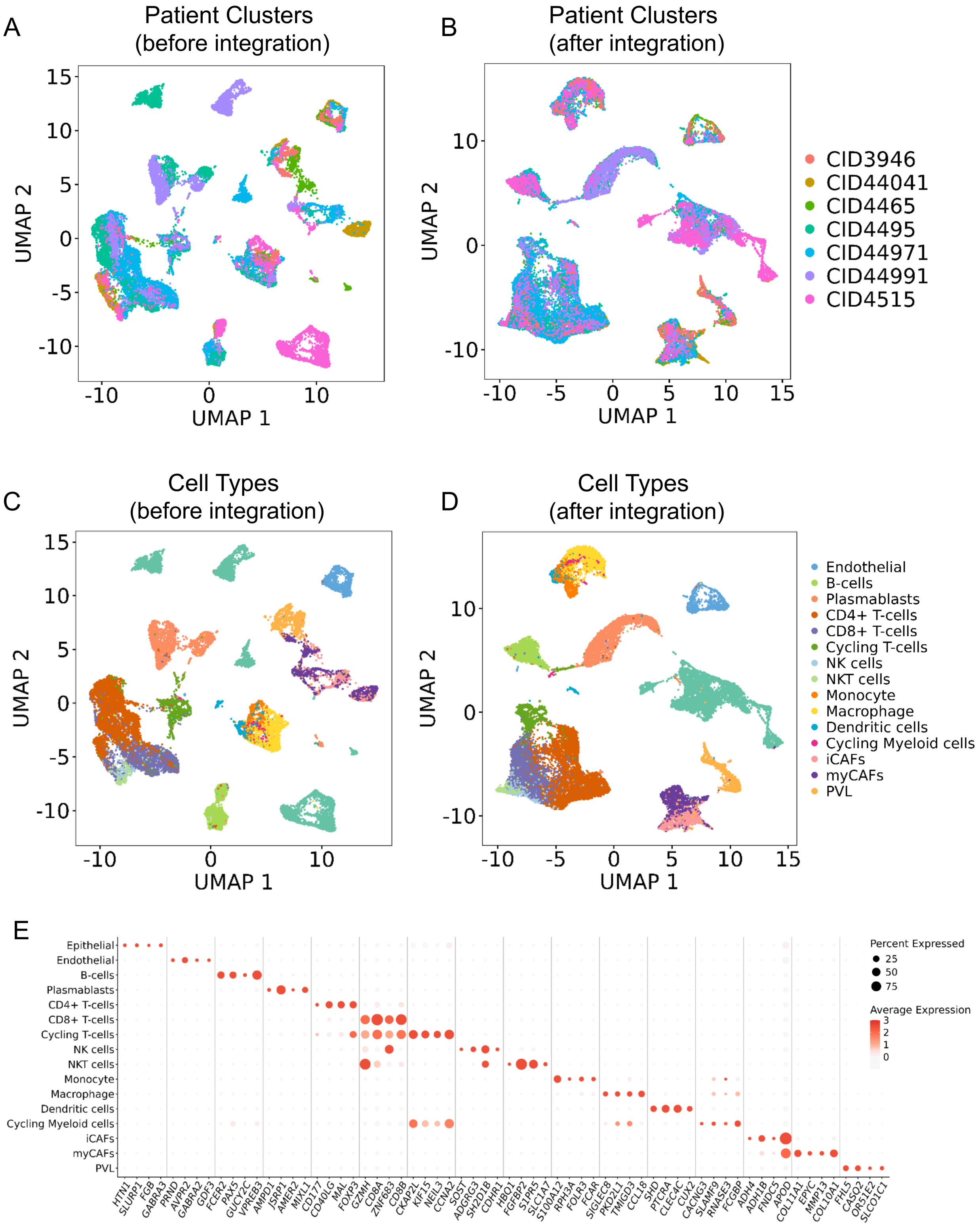


**Supplementary Figure S13. Batch integration and cell type-specific marker expression in the TNBC single-cell dataset.**

A) UMAP visualizations showing patient clusters before and after batch integration. B) UMAPs showing cell type clustering before and after batch integration. C) Dot plot illustrating the expression pattern of top four marker genes for each study-reported cell type cluster.

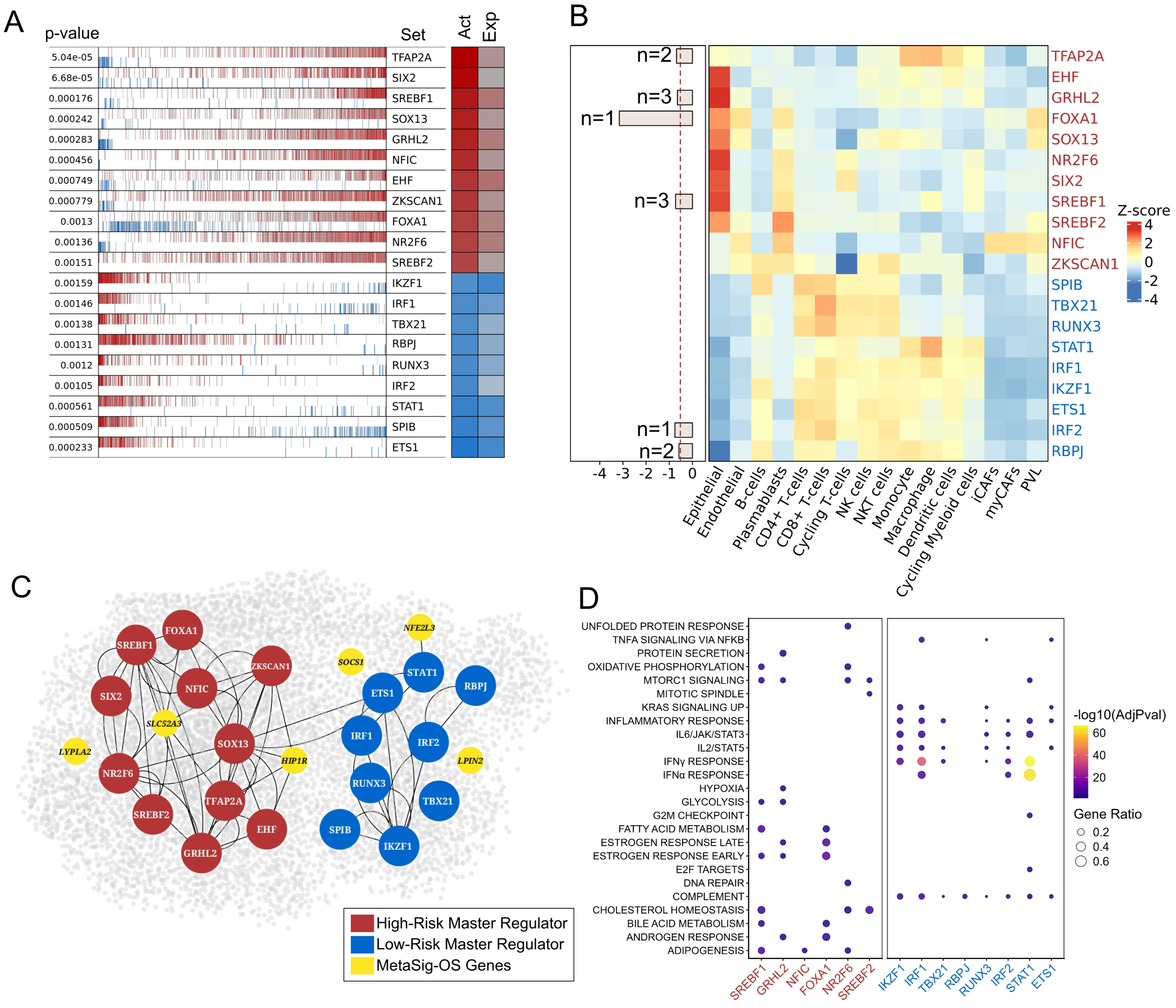


**Supplementary Figure S14. Upstream Regulatory Architecture of MetaSig-OS.** A) Plot showing the top 20 activated and repressed master regulators (MRs) identified between high- and low-risk patients using VIPER analysis. The leftmost column indicates the significance (p-value) of each regulon. The central barcode-like panel represents the distribution of activated (red) and repressed (blue) target genes ranked according to the differential expression signature (left: most down-regulated; right: most up-regulated). The “Act” column represents inferred regulon activity, while the “Exp” column indicates the expression level of the corresponding transcription factor. B) Heatmap showing the mean scaled AUCell regulon activity scores of the MetaSig-OS master regulators across cell types in the TNBC single-cell dataset. Colors represent the mean scaled AUCell score for each regulon within each cell type. The horizontal bar plot summaries gene essentiality based on DepMap dependency scores across 19 TNBC cell lines. Bar length corresponds to the mean dependency score, while *n* indicates the number of TNBC cell lines in which the corresponding master regulator was classified as essential. C) Network representation of MRs and their association with MetaSig-OS genes. Red nodes indicate MRs activated in high-risk patients, whereas blue nodes represent MRs activated in low-risk patients. Yellow nodes denote MetaSig-OS genes connected to the corresponding regulatory network. Edges represent inferred regulatory interactions between transcription factors and target genes. D) Dot plot depicting pathways represented by high- and low-risk patient-specific MRs. Dot size represents gene ratio, and color intensity indicates significance (-log10 adjusted p-value).

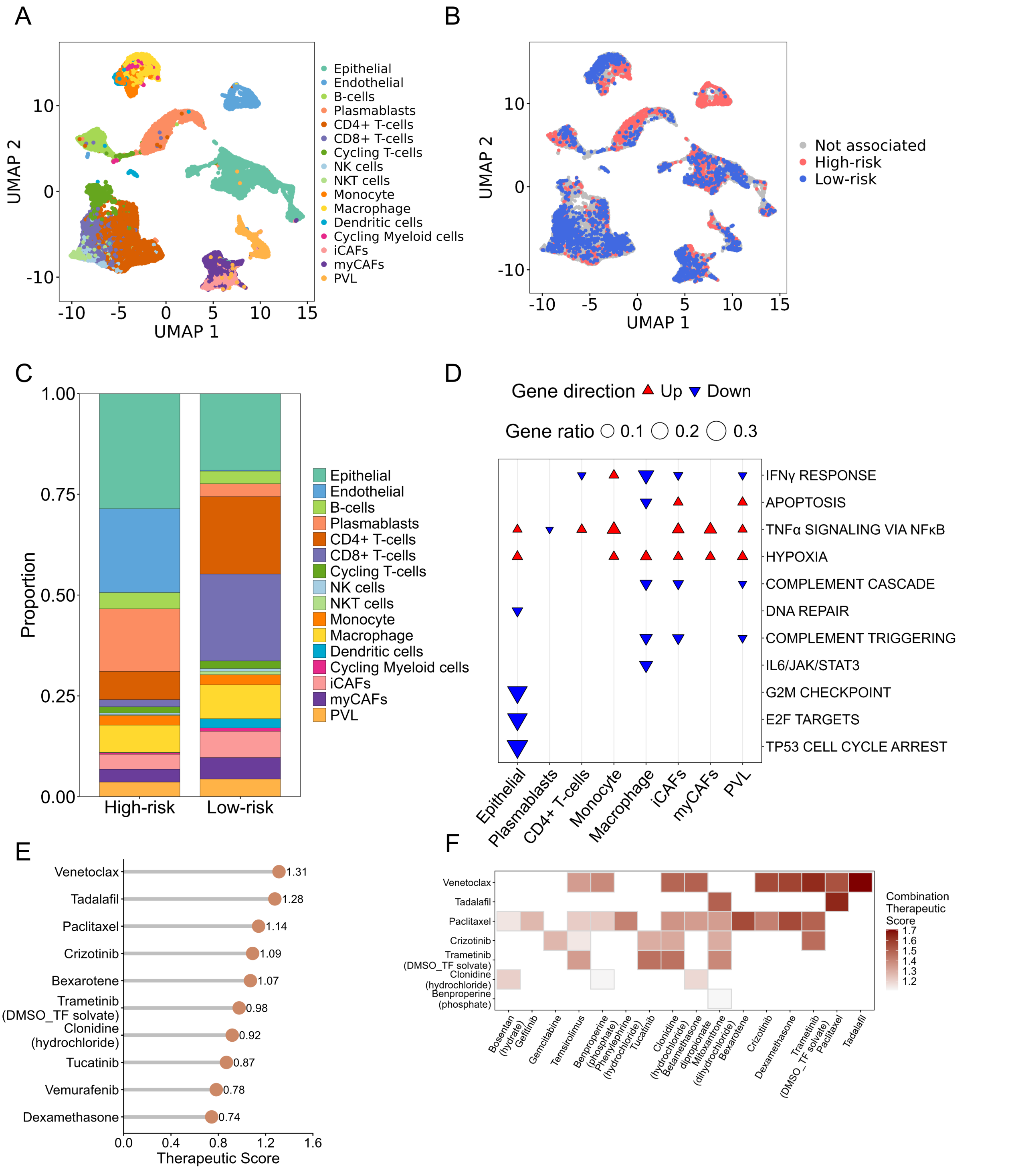


**Supplementary Figure S15. Risk-Associated Cell States and Candidate Therapeutics for MetaSig-OS.** A) UMAP visualization showing the distribution of annotated cell types in the TNBC single-cell dataset. B) UMAP depicting the distribution of cells associated with high-risk and low-risk patients across the single-cell landscape. Red cells represent populations associated with high-risk patients, blue cells represent populations associated with low-risk patients, and grey cells indicate populations not significantly associated with either risk group. C) Stacked bar plot showing the proportional distribution of annotated cell types among cells associated with high-risk and low-risk patients. D) Pathway analysis comparing high-risk and low-risk associated cells across individual cell types. Red upward triangles indicate pathways significantly up-regulated in high-risk cells, whereas blue downward triangles represent pathways enriched in low-risk cells. Dot size corresponds to gene ratio, reflecting the proportion of pathway-associated genes contributing to the signal. E) Top FDA-approved therapeutic drugs identified through drug repurposing analysis using differential gene expression signatures derived from epithelial cells associated with high- versus low-risk patients in the Tahoe perturbation dataset. F) Heatmap showing significant candidate drug combinations predicted for high-risk patients. Color intensity represents the combination therapeutic score, with darker shades indicating stronger predicted synergistic therapeutic potential.

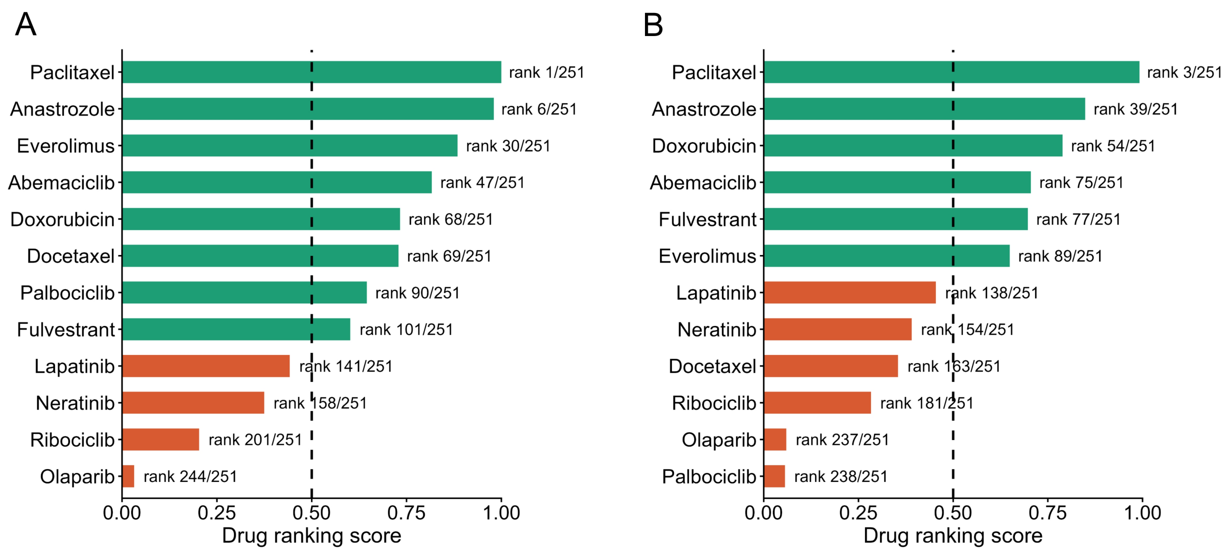


**Supplementary Figure S16. Positive Control Validation for Drug Repurposing.** A) Positive control validation plot for MetaSig-EFS. B) Positive control validation plot for MetaSig-OS.
